# RNase L reprograms translation by widespread mRNA turnover escaped by antiviral mRNAs

**DOI:** 10.1101/486530

**Authors:** James M Burke, Stephanie L Moon, Evan T Lester, Tyler Matheny, Roy Parker

## Abstract

In response to foreign and endogenous double-stranded RNA (dsRNA), protein kinase R (PKR) and ribonuclease L (RNase L) reprogram translation in mammalian cells. PKR inhibits translation initiation through eIF2α phosphorylation, which triggers stress granule (SG) formation and promotes translation of stress responsive mRNAs. The mechanisms of RNase L-driven translation repression, its contribution to SG assembly, and its regulation of dsRNA stress-induced mRNAs are unknown. We demonstrate that RNase L drives translational shut-off in response to dsRNA by promoting widespread turnover of mRNAs. This alters stress granule assembly and reprograms translation by only allowing for the translation of mRNAs resistant to RNase L degradation, including numerous antiviral mRNAs such as *IFN-β*. Individual cells differentially activate dsRNA responses revealing variation that can affect cellular outcomes. This identifies bulk mRNA degradation and the resistance of antiviral mRNAs as the mechanism by which RNaseL reprograms translation in response to dsRNA.

## INTRODUCTION

Double-stranded RNA (dsRNA) is a pathogen-associated molecular pattern (PAMP) generated during viral infection that can initiate the innate immune response (Jensen and Thomsen, 2012). Endogenous “self’ dsRNAs can also initiate the innate immune response, and dysregulation of cellular pathways that reduce self-dsRNA causes human diseases, such as Aicardi–Goutières syndrome (AGS) (Pestal et al., 2015, Liddicoat et al., 2015, George et al., 2016, Li et al., 2017). Elevated levels of endogenous dsRNAs have also been implicated in contributing to neurodegenerative diseases, such as ALS (Saldi et al., 2014, Krug et al. 2017), and can contribute to chronic inflammation associated with cancers and autoimmune disorders (Grivennikov et al., 2010, Waldner 2009). Nevertheless, the mechanisms by which mammalian cells respond to dsRNA remain incompletely understood.

Several pattern recognition receptors (PRRs) recognize dsRNA in mammalian cells (Jensen and Thomsen, 2012). Recognition of dsRNA by RIG-I, MDA-5, or TLR3 activates the transcription factors IRF3 and/or IRF7 thereby inducing type-1 interferons (IFNs) and inflammatory cytokines, which prime the antiviral state of cells via autocrine and paracrine JAK/STAT signaling and promote cell-mediated innate and adaptive immune responses (Ivashki et al., 2014). Concurrent with the induction of antiviral genes, global translation is reduced in response to dsRNA by protein kinase R (PKR) and ribonuclease L (RNase L) as part of a process termed host shutoff. Host shutoff promotes an antiviral cellular state by limiting viral gene expression and transforming the functional cellular transcriptome (Iordanov et al., 2000).

Activation of PKR by dsRNA results in phosphorylation of eIF2α on serine 51, which reduces canonical translation initiation and promotes the translation of stress response mRNAs that use non-canonical translation initiation (Dalet et al., 2015). This also triggers the formation of stress granules (SGs), conserved RNA-protein complexes that contain non-translating mRNAs, RNA-binding proteins – G3BP1, PABPC1, TIA1 – and several key antiviral PRRs – OAS/RNase L, PKR, MDA-5, and RIG-I (García et al., 2006, Onomoto et al., 2012, Reineke et al., 2012, Yoo et al., 2014). Many viruses inhibit SG assembly by diverse means (reviewed in Lloyd, 2013), suggesting that SGs serve as antiviral signaling hubs and/or reduce viral replication through the sequestration of viral mRNAs/proteins (Buchan and Parker, 2009; Kang et al., 2018). However, the disassembly of SGs via dephosphorylation of p-eIF2α by GADD34, which is induced by IRF3, has been proposed to promote translation of stress-induced antiviral mRNAs that are sequestered to SGs (Kojima et al., 2003, Dalet et al., 2017). Thus, the mechanisms and functions of SG assembly/disassembly during the dsRNA/antiviral response remain unclear.

RNase L is an endonuclease activated by oligo(2’-5’A), which is produced when OAS proteins bind to dsRNA. RNase L is thought to act in a non-specific manner to cleave ssRNA regions in Y-RNAs, tRNAs, rRNAs, and host/viral mRNAs (Clemens and Williams, 1978, Andersen et al., 2009, Chakrabarti et al., 2011, Brennan-Laun et al., 2014, Donovan et al., 2017). These activities of RNase L reduce viral gene expression and replication by arresting global translational and promoting apoptosis (Zhou et al., 1997). RNase L is proposed to arrest translation by either cleavage of rRNA or production of RNA cleavage fragments that signal for translational arrest (Wreschner et al., 1981; Donovan et al., 2017). However, because these modes of translational arrest are presumably non-specific, a long-standing mystery in the field is how dsRNA-induced antiviral mRNAs would be translated during RNase L-driven translational arrest.

We present data demonstrating that RNase L promotes widespread degradation of cellular mRNAs in response to dsRNA, which leads to a decrease in bulk translation. This process alters stress granule assembly and promotes PABPC1 translocation from the cytosol to the nucleus. Strikingly, mRNAs encoding key antiviral and inflammatory cytokines, such as the *IFN-β* and *IL-6* mRNAs, escape RNase-L mediated mRNA turnover, which permits their translation during host shut-off when bulk mRNA turnover is the primary driver of global translation repression.

## RESULTS

### RNase L catalytic activity alters SG assembly during the dsRNA response

RNase L represses translation and accumulates in stress granules (Onomoto et al., 2012, Reineke et al., 2015). Thus, we examined if RNase L activity affected stress granule assembly. We generated RNase L knockout (RL-KO) A549 and U-2 OS cell lines using CRISPR-Cas9, and then reconstituted expression of either RNase L or catalytically inactive RNase L-R667A in the RL-KO cells via lentiviral transduction or transient transfection (Figure S1A,B,C). Cells were transfected with high molecular weight poly(I:C), a viral dsRNA mimic that induces PKR-dependent SG assembly and activates the OAS/RNase L pathway. SG assembly was assessed by immunofluorescence assay (IF) for SG-associated proteins G3BP1 and PABPC1.

In comparison to the parental (WT) cell lines, we observed two distinct phenotypes in RL-KO cell lines that were rescued by expression of RNase L, but not RNase L-R667A. First, SGs in the RL-KO cells were canonical in morphology (large and irregular in shape), whereas cytoplasmic puncta of G3BP1 and PABPC1 observed in the WT cells were invariably small and punctate (Figures 1A,B,C and S1D,E). Second, a substantial fraction of PABPC1 translocated from the cytosol to the nucleus in WT cells, whereas PABPC1 remained localized in the cytosol and SGs in RL-KO cells. The RNase L-dependent reduction in SG size was specific to the dsRNA stress response, as sodium arsenite treatment induced canonical SGs in both WT and RL-KO cells (Figure S1F).

**Figure 1.**
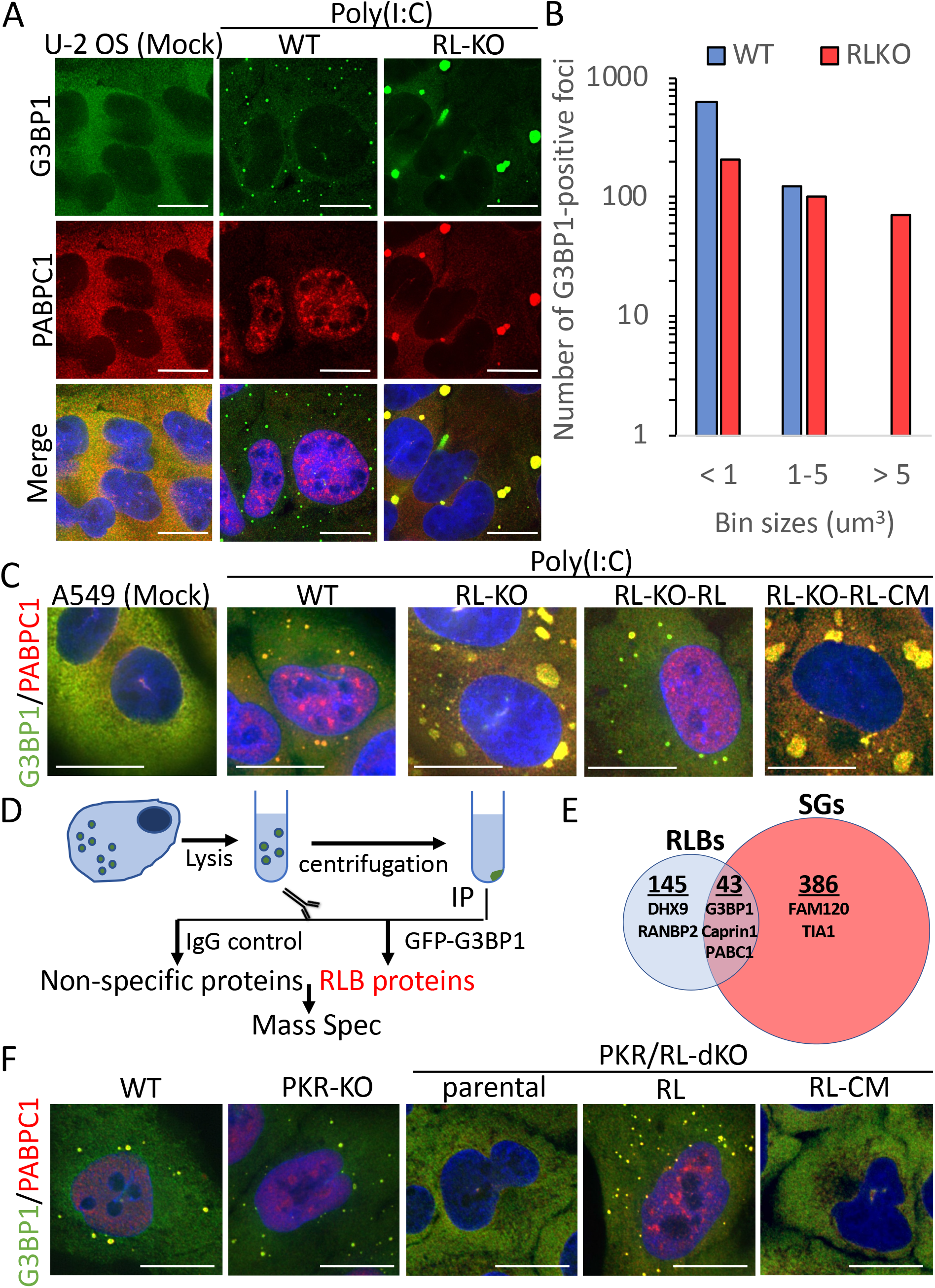
RNase L catalytic activity limits SG assembly and promotes the assembly of compositionally distinct RNP bodies. (A) IF for SG-associated proteins G3BP1 and PABPC1 in WT and RL-KO U-2 OS cells eight-hours post-transfection of poly(I:C) (500-ng/ml). Nuclei were stained with DAPI (blue). (B) G3BP1-positive foci binned by volume in WT and RL-KO from greater than 30 cells in 3 fields of U-2 OS cells. (C) IF for G3BP1 and PABPC1 in WT and RL-KO A549 cells 8 hours post-poly(I:C). Images for G3BP1 and PABPC1 staining are shown in Figure S1D. (D) Schematic of approach for RLBs purification. (E) Proteins identified by RLB mass spectrometry analysis and overlap with sodium arsenite-induced SG proteome (Data File S1). (F) G3BP1 and PABPC1 IF in PKR-KO and PKR and RNase L double KO (PKR/RL-KO) A549 cells rescued with RNase L (RL) or RNase L-R667A (RL-CM) six hours post-poly(I:C). Scale bars represent 15 μm.

Several observations indicate that the small punctate foci of PABPC1 and G3BP1 in WT cells, which we refer to as RNase L-dependent bodies (RLBs), are distinct from SGs. First, IF revealed that while RLBs share several proteins with dsRNA-induced SGs in RL-KO cells (G3BP1, Caprin1, Ataxin-2, PABPC1), multiple common SG components are not enriched within RLBs, including TIA, FAM120A, PUM1, and FXR1 (Figure S2A,B). Mass spectroscopy of purified RLBs identified 188 proteins associated with RLBs that only partially overlap with the SG proteome (Figures 1D,E and S2C and Data File S1). Second, although RLBs contain poly(A)+ mRNAs as assessed by FISH (Figure S3A), they form in the presence of cycloheximide (Figure S3B), which traps mRNAs in polysomes and blocks SG formation (Protter and Parker, 2016). Third, RLBs are not formed via p-eIF2α-mediated translation repression since they form in the presence of ISRIB (Figure S3B), which prevents p-eIF2α-driven translational repression and SG assembly (Sidrauski et al., 2015). Fourth, RLBs form independently of PKR, as PKR-KO cells produced RLBs in response to poly(I:C) (Figures 1F and S3C,D,E,F). Finally, RLBs require RNase L activity, as RLBs were absent in PKR/RNase L double-KO cells in response to poly(I:C), and reconstitution of RNase L but not RNase L-R667A restored their assembly (Figures 1F and S3C-F). Thus, RNase L activation inhibits canonical SG assembly and promotes the assembly of RLBs.

### RNase L initiates rapid and widespread turnover of mRNAs

Since mRNAs are an integral component of SGs (Van Treek et al., 2018), and PABPC1 translocation to the nucleus occurs upon mRNA turnover (Glaunsinger and Ganem, 2004; Kumar et al., 2011), we hypothesized that RNase L limits SG formation by degrading SG mRNAs. Thus, we examined whether RNase L alters the localization and/or abundance of mRNAs that enrich within SGs by singlemolecule fluorescent in situ hybridization (smFISH). Strikingly, at two- and six-hours post-transfection of poly(I:C), the SG-enriched *AHNAK* mRNA (Khong et al., 2017) was strongly reduced in the cytosol and did not localize to RLBs in WT cells (Figure 2A,B). In contrast, *AHNAK* mRNA remained abundant and localized to SGs in RL-KO cells. qRT-PCR analysis confirmed these results, showing that *AHNAK* mRNA levels significantly decreased in WT, but not RL-KO cells, post-poly(I:C) (Figure 2C). Analysis of the SG-enriched NORAD long non-coding RNA (lncRNA) yielded similar results (Figure S3G,H,I). These data suggest that RNase L promotes degradation of RNAs that localize to SGs.

**Figure 2.**
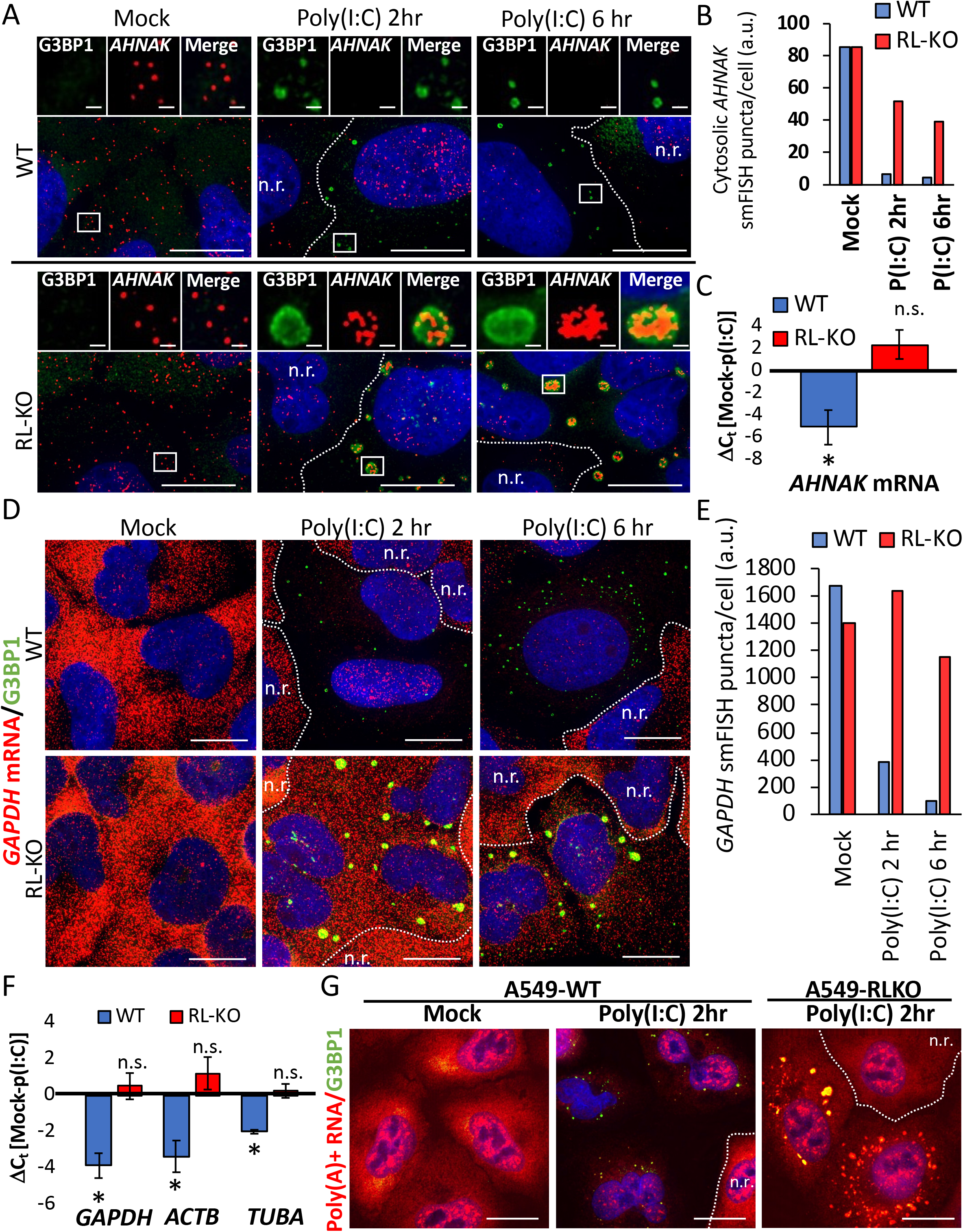
RNase L promotes widespread turnover of mRNAs. (A) smFISH for *AHNAK* mRNA in WT and RL-KO U-2 OS cells +/− poly(I:C) (500-ng/ml) for two or six hours with G3BP1 as a RLBs/SG marker. N.r. indicates non-responsive cells with respect to RLB/SG assembly. (B) Quantification of *AHNAK* mRNA smFISH from (A), with analysis of 3 fields of 17-30 RLB/SG cells. (C) RT-qPCR analysis of AHNAK mRNA in WT and RL-KO A549 cells at zero- and six-hours post-poly(I:C) transfection. (D) smFISH for *GAPDH* mRNA (red) in WT and RL-KO U-2 OS cells two hours post-poly(I:C). (E) Quantification of *GAPDH* mRNA smFISH. (F) qRT-PCR quantification of *GAPDH, Actin B,* and *Tubulin A* mRNAs in WT and RL-KO A549 cells transfected with or without (Mock) poly(I:C) for six hours. Bars represent the average Ct value +/−S.E.M. from at least five independent experiments. (G) Oligo(dT) (red) and G3BP1 (green) staining of WT and RL-KO A549 cells two hours post-poly(I:C). Quantification of mean signal in SG+ cells shown in Figure S4J. Scale bars represent 15 μm.

We also examined whether RNase L-mediated mRNA turnover affected mRNAs not enriched in SGs. We observed that the *GAPDH* mRNA, which is abundant and depleted from SGs (Khong et al., 2017), was reduced by at least 90% in WT cells containing RLBs by two hours post-poly(I:C) (Figure 2D,E). In neighboring cells that lack RLBs and thus were presumably not transfected with poly(I:C) and lack RNase L activity, *GAPDH* mRNA levels remained abundant and comparable to mock-treated cells. Importantly, *GAPDH* mRNA levels were unchanged in response to poly(I:C) in RL-KO cells, indicating RNase L is required for the reduction in *GAPDH* mRNA levels. qRT-PCR analysis confirmed these observations, revealing that *GAPDH* mRNA levels, as well as *Actin B* and *Tubulin A* mRNAs, decreased (greater than 75%) in WT but not RL-KO cells following poly(I:C) transfection (Figure 2F). Moreover, FISH for poly(A)+ RNAs revealed at least a 70% RNase L-dependent decrease in cytosolic poly(A)+ RNAs by two hours post-poly(I:C) in cells containing RLBs (Figures 2G and S3J). Taken together, these results suggest RNase L is degrading the majority of cytoplasmic mRNAs in response to dsRNA.

### *IFN-β* and *IL-6* mRNAs escape RNase L-mediated mRNA turnover

The widespread degradation of host mRNAs by RNase L creates a problem for how cells undergoing the antiviral/dsRNA response are able to produce proteins from IRF3-induced genes, such as IFN-β. One possibility is that IRF3-induced mRNAs escape RNase L-dependent turnover. To test this, we performed smFISH for the *IFN-β* mRNA simultaneously with the *GAPDH* mRNA to determine if *IFN-β* mRNAs were present in cells that have degraded the *GAPDH* mRNA. Strikingly, in WT cells wherein *GAPDH* mRNA had been reduced by RNase L in response to poly(I:C), cytosolic *IFN-β* mRNA was abundant (Figures 3A and S4A). Moreover, RT-qPCR revealed that the *IFN-β* mRNA was induced to comparable levels between WT and RL-KO cells six hours post-poly(I:C) (Figure 3B). These data indicate that the *IFN-β* mRNA is resistant to RNase L-mediated degradation.

**Figure 3.**
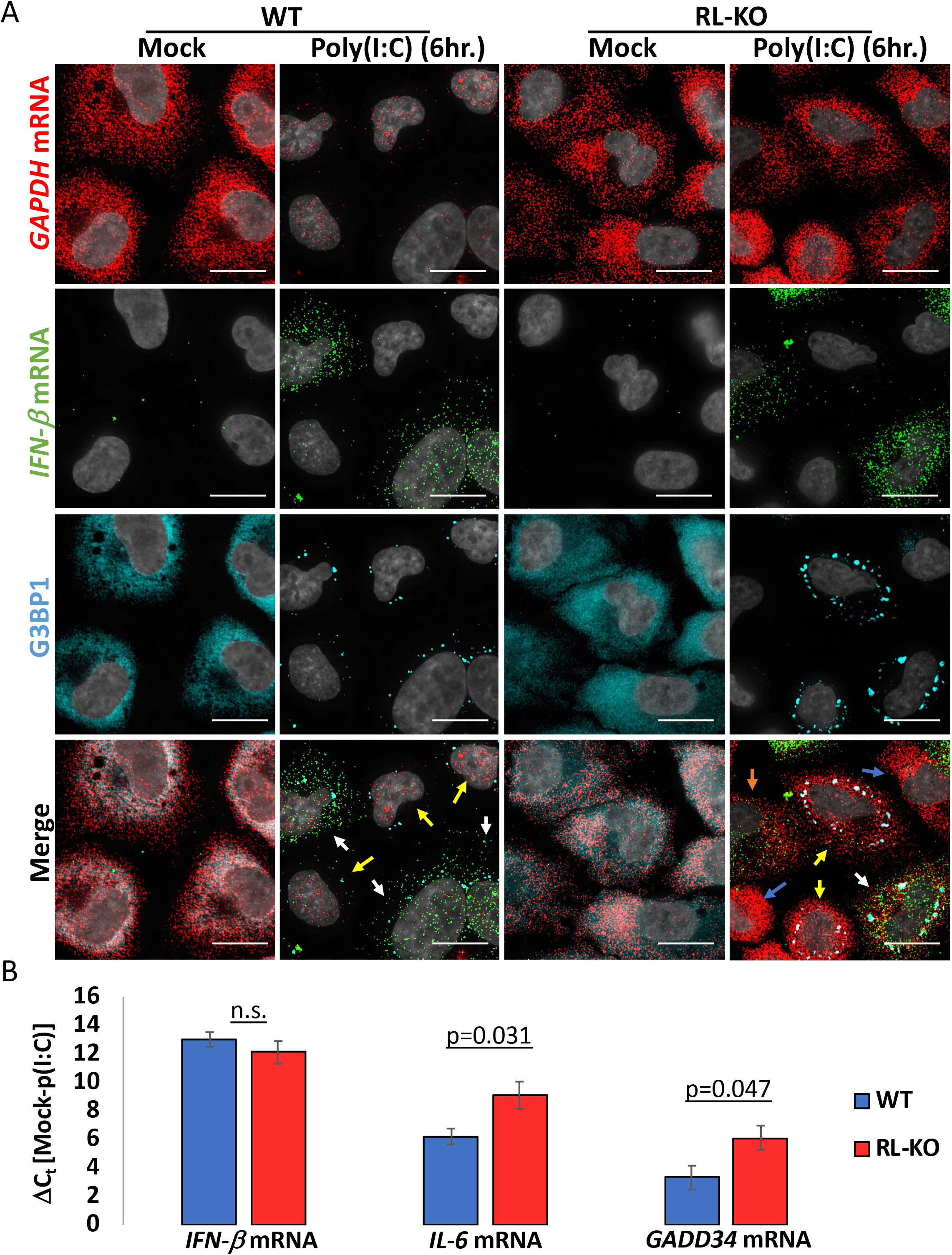
*IFN-β* mRNA escapes RNase L-mediated mRNA turnover. (A) smFISH/IF for *IFN-β* mRNA (green) *GAPDH* mRNA (red) and IF for G3BP1 (cyan). Nuclei were stained with DAPI (gray). WT and RL-KO A549 cells +/− poly(I:C) with smFISH/IF performed six hours post-transfection. Cells that contain RLBs/SGs and *IFN-β* mRNA indicated by white arrows. Cells with RLBs/SGs without *IFN-β* mRNA are indicated by yellow arrows. Cells with *IFN-β* mRNA without RLBs/SGs are indicated by orange arrows. Cells lacking both RLBs/SGs and *IFN-β* mRNA indicated by blue arrows. Scale bars represent 15 μm. (B) RT-qPCR analysis of *IFN-β, IL-6,* and *GADD34* mRNA expression in WT and RL-KO A549 cells six-hours post-transfection of poly(I:C). Bars represent the average Ct value +/− the S.E.M from greater than 5 independent experiments.

We observed a similar phenomenon assaying the *IL-6* mRNA via smFISH (Figure S4B), although there were more IL-6 mRNAs present in the RL-KO cells, which suggests the IL-6 mRNA is only partially resistant to RNase L degradation. Consistent with this, qRT-PCR revealed that *IL-6* mRNA levels were ~8-fold lower in WT cells as compared to RL-KO cells post-poly(I:C) (Figure 3B). Similarly, the *GADD34* mRNA is induced by poly(I:C), but is reduced in an RNase L-dependent manner as assessed by qRT-PCR. Thus, there are mRNA specific differences in resistance to RNase L degradation, even among mRNAs induced by IRF3/7.

### Differential PRR activation in individual cells in response to dsRNA

Our single-cell analyses of mRNAs, RLBs, and SGs revealed a remarkable cell-to-cell variability in the response to dsRNA. For example, while the six WT cells treated with poly(I:C) in Figure 3A all showed RNase L activation, as determined by the formation of RLBs and degradation of GAPDH mRNA, only three cells strongly induced the *IFN-β*mRNA (Figure 3A, white arrows). We also observed cells that induced the *IFN-β* mRNA without activation of RNase L (Figure S4C). Similarly, in RL-KO cells, we can observe individual cells that activate PKR (as assessed by SG formation), with or without induction of the *IFN-β* mRNA (Figure 3, white and yellow arrows). We also observed RL-KO cells that induced the *IFN-β* mRNA without activation of PKR (Figure 3A, orange arrow). Interestingly, WT cells only generate RLBs and not SGs (Figure 1), indicating that PKR activation does not commonly occur without co-activation of RNase L in individual WT cells. These differences are not due to failures in transfections since cells are activating one portion of the response to dsRNA, but not other pathways. This demonstrates that individual cells respond differentially to dsRNA, which can allow multiple cellular outcomes during the dsRNA/antiviral response (see discussion).

### Genome-wide analysis of RNase L-mediated mRNA degradation

To identify the RNase L sensitive and RNase L resistant mRNAs on a comprehensive scale, we performed high-throughput RNA sequencing on WT and RL-KO A549 cells before and after poly(I:C) transfection. Standard differential expression analysis showed that few RNAs were significantly different between untreated WT and RL-KO cell lines (Figure S5A). In response to poly(I:C), a substantial number of RNAs significantly decreased or increased in WT cells (Figure 4A). In contrast, only a small number of RNAs significantly increased in RL-KO cells. We then normalized RNA levels to ERCC spike-in control RNAs to control for the expected gross reductions in mRNA. This was a necessary step since the number of upregulated and downregulated RNAs in WT cells post-poly(I:C), as well as the magnitude by which they changed, was overestimated and underestimated, respectively, by standard differential expression analyses (Figures S5B,C,D,E). We then calculated the differential in ERCC-normalized RNA levels between WT and RL-KO cells in response to poly(I:C) and plotted as a color gradient on the scatterplot of WT cells treated with or without poly(I:C) (Figure 4B). These analyses revealed several important observations.

**Figure 4.**
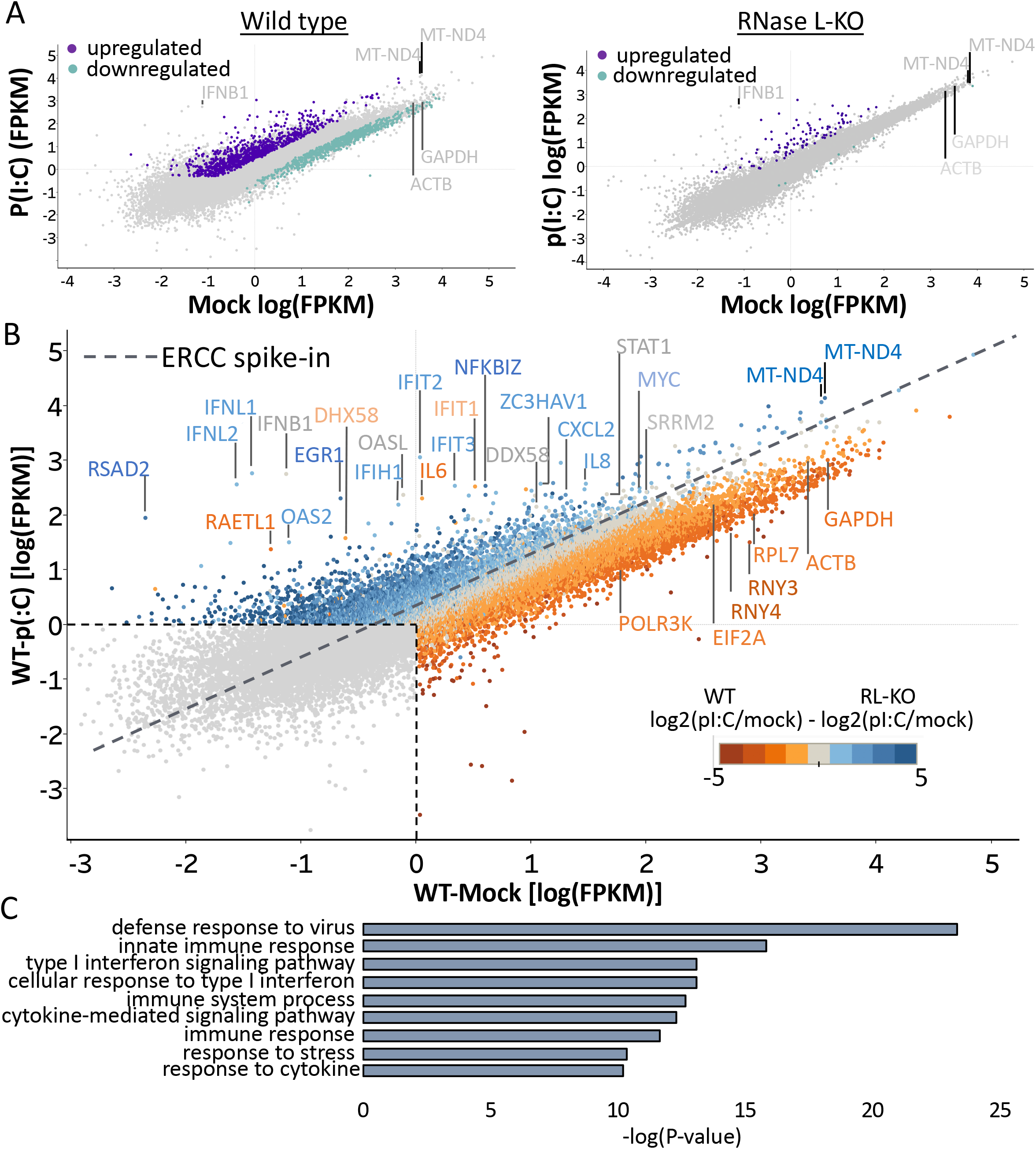
RNase L regulates global mRNA levels during dsRNA stress. (A) Differential expression analysis without normalization from total RNA isolated from WT or RNase L-KO A549 cells before and after poly(I:C) transfection. (Upregulated transcripts = Fold Change > 2, p-value < 0.05; downregulated transcripts = Fold Change < 0.5, p-value < 0.05). (B) Differential expression plot of mRNAs in WT cells treated with or without poly(I:C). The linear regression of the ERCC spike-in control RNAs is represented by dashed line. The color of the dots indicates their ERCC-normalized differential expression relative to RL-KO cells transfected +/− poly(I:C). Orange = RNase L-dependent decrease, blue = RNase L-dependent increase. (C) GO analysis of mRNAs that are induced in both WT and RL-KO cells post-poly(I:C) (greater than 4-fold, FPKM > 1 in either WT or RL-KO cells with poly(I:C) treatment) from Data file S1.

First, we observed that poly(I:C) transfection in WT cells led to a striking reduction in essentially all abundant mRNAs, with RNA transcripts from 6,310 genes being reduced by 2-fold or more (Figure 4B and Data File S1). This was most notable for abundant mRNAs with long steady-state half-lives (Figure S6A,B,C,D,E), which is consistent with our qRT-PCR analysis of *GAPDH, Actin B,* and *Tubulin A* mRNAs (Figure 2F). Importantly, the decrease in these mRNAs is largely due to RNase L since the vast majority of abundant mRNAs did not substantially change in level in response to poly(I:C) in RL-KO cells (Figure S5B,D). Mitochondrial mRNAs (i.e. MT-ND4 and MT-ND4L) were unaffected, and in fact slightly increased in WT cells relative to RL-KO cells post-poly(I:C), effectively serving as an internal control that suggests our analyses may underestimate the level of RNase L-dependent mRNA reductions (Figures 4B and S5D,E). These results confirm that RNase L promotes the degradation of the majority of abundant and stable cytoplasmic mRNAs.

Second, we identified a population of mRNAs that substantially increases in both WT and RL-KO cells in response to poly(I:C) (Figures 4B and S5D,E and Data File S1). This population of mRNAs, which includes *IFN-β*, *IFIT2, OAS2, IFIH1 (MDA5), DDx58 (RIG-I)* and *IL-6,* is highly enriched for IRF3-induced mRNAs and interferon-stimulated genes (ISGs) (Figure 4C). This is consistent with our smFISH and RT-qPCR analyses and suggests that dsRNA-induced antiviral mRNAs escape RNase L-mediated mRNA turnover (Figure 3). Some mRNAs, such as the *IL-6* and *RAET1L* mRNAs, are induced to higher levels in the RL-KO cell line suggesting they are partially degraded by RNase L. Assuming transcriptional activation of these mRNAs is similar in the two cells lines, these data suggest there is a range of RNase L resistance amongst dsRNA-induced mRNAs.

The dsRNA-induced mRNAs resistant to RNase L do not have substantial differences in GC-content, 5’ UTR length, 3’ UTR length, and total transcript length of in comparison to RNase L sensitive mRNAs (Data File S1), suggesting that these features may not contribute to RNase L resistance. MEME analysis of their 3’ UTRs did not identify a notable sequence motif common to all of these mRNAs, though many of these mRNAs (31%) contain AU-rich elements (AREs) (Data File S1), a common motif in mRNAs encoding cytokines (Savan, 2014). Meta-analysis of CLIP studies for RBPs did not reveal a significant enrichment in association with RBPs and these mRNAs (data not shown). However, we note that many of these mRNAs are expressed at very low levels without induction by dsRNA, and the CLIP studies were not performed during dsRNA stress. Therefore, the mechanisms by which these mRNA escape RNase L-mediated mRNA turnover remains undefined.

Third, we identified constitutively expressed mRNAs (i.e. *MYC, SRRM2, STAT1)* that do not decrease in WT cells post-poly(I:C). Interestingly, STAT1 mediates interferon signaling for ISG production, and MYC is a negative regulator of IRF7 (Kim et al., 2016). We note that *STAT1* has been shown to be induced by type I and type II interferons in some studies (Rusinova et al., 2013), and thus we cannot rule out that it is sensitive to RNase L, but also transcriptionally induced. Nevertheless, these data suggest that a portion of constitutively expressed mRNAs, some of which regulate the antiviral response, also escape RNase L-mediated mRNA turnover.

Fourth, we identified a population of low abundance mRNAs whose levels increase specifically in WT cells post-poly(I:C) (Figure 4B). We note that several non-mutually exclusive possibilities may account for this. First, despite our use of the ERCC spike-in control, this may be a technical artifact of high-throughput sequencing, whereby reads of RNAs that are less efficiently turned over by RNase L are artificially inflated due to the reduction in the majority of abundant mRNAs. Second, mRNAs that already have rapid decay rates may be not affected by RNase L since the overall decay rate is not changed significantly (Figure S6E). Third, RNase L activation may stabilize a subset of mRNAs with short halflives. Finally, RNase L activation may promote the expression of numerous genes.

Combined, these data confirm that RNase L activation promotes the turnover of abundant mRNAs in response to poly(I:C), whereas highly-induced antiviral mRNAs are resistant to this process.

### RNase L drives translational repression via bulk mRNA turnover

The rapid and widespread RNase L-dependent turnover of mRNAs led us to examine if this contributes to dsRNA-induced translational repression. Currently, RNase L is thought to repress translation by degrading rRNA (Wreschner et al., 1981), or by RNase L-cleaved RNAs inhibiting translation (Donovan et al., 2017), possibly by triggering phosphorylation of eIF2 *a* by PKR, which is the primary eIF2 *a* kinase activated by dsRNA (Dalet et al., 2015). To resolve the relative contributions of these mechanisms to RNase L-driven translational repression, we evaluated dsRNA-induced translational repression in cell lines with or without RNase L and/or PKR.

Our data argue that RNase L activity drives rapid translation repression independently of PKR in the majority of A549 cells in response to dsRNA. The key observation is that metabolic labeling of nascent proteins with either ^35^S-labeled methionine and cysteine or puromycin revealed that both WT and PKR-KO cells strongly repressed translation by two-hours post-poly(I:C), whereas RL-KO cells maintained translation (Figures 5A,B and S7A,B).

**Figure 5.**
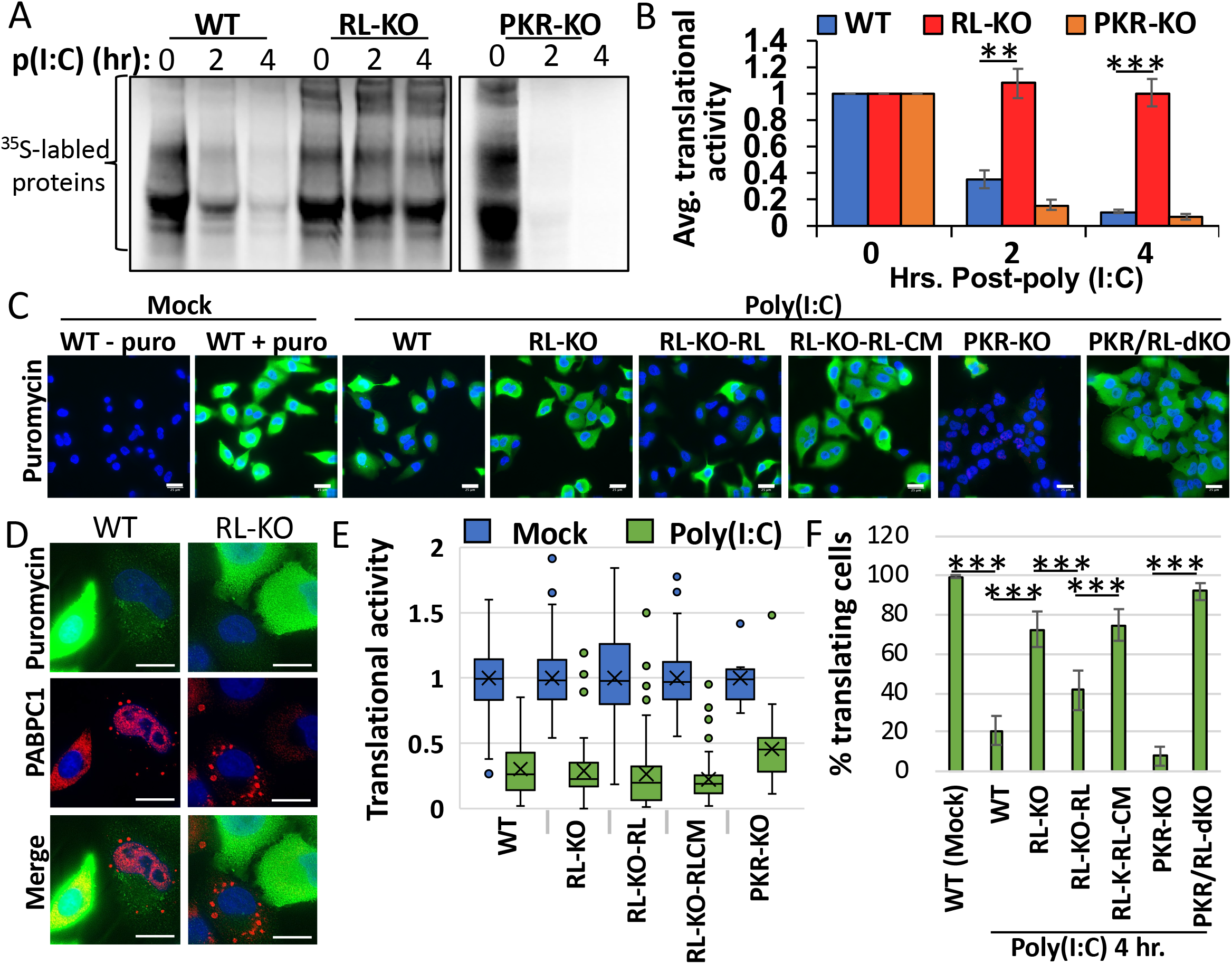
RNase L drives translational shut-off. (A) S-35 metabolic labeling of newly synthesized proteins in WT, RL-KO, and PKR-KO A549 cells post-poly(I:C). (B) Quantification of S-35-labeling experiments as represented in (A). Bars represent the average signal normalized to mock treatment from at least two independent replicates +/− S.D. normalized to time zero. (C) Representative images from SUnSET puromycin-labeling (green) analysis of indicated cells lines four hours post-poly(I:C) transfection. Dapi-stained nuclei (blue). Scale bar is 25 μm. (D) Similar to (C) but enlarged to show correlation between RLBs/SGs via PABPC1 IF (red) and translational activity via puromycin IF (green). Scale bar is 15 μm. (E) Quantification of the mean intensity normalized to mock-treated cells of puromycin staining in cells (between 51-176 cells analyzed for each cell line) that contain RLBs/SGs four hours post-poly(I:C). (F) Quantification of the percentage of cells translating (puromycin signal greater than 80% of average signal from untreated cells) four hours post-poly(I:C). Bars represent the average +/− S.D. normalized to untreated cells from at least three independent replicates in which cells from at least five fields of view were analyzed. * indicates p-value <0.05, ** indicates p-value < 0.005, and *** indicates p-value <0.001 as determined by student’s t-test.

The lack of translational repression in the RL-KO cells was surprising since some cells generate PKR-dependent SGs (Figure 1), which we hypothesized might be due to variation in the response of individual cells. Therefore, we examined translation in individual cells by IF for puromycin-labeled proteins (Schmidt et al., 2009). This analysis revealed that the RL-KO cells containing SGs in fact repressed bulk translation in response to poly(I:C), which was dependent on PKR, as PKR/RL-dKO cells did not repress translation in response to poly(I:C) (Figure 5C,D,E,F). However, less than 30% of the RL-KO cells repressed translation in response to poly(I:C) (Figure 5F). In contrast, greater than 80% of the WT and PKR-KO cells underwent translational repression and showed RLB body formation with nuclear translocation of PABPC1, hallmarks of RNase L activation (Figure 5C,D,E,F). Thus, RNase L represses translation in the majority of A549 cells in response to dsRNA, although in the absence of RNase L, PKR can repress translation in a minority of cells, providing additional evidence for variation in how individual cells respond to dsRNA.

In addition to degrading mRNAs, RNase L could repress translation by degrading rRNA or by activating signaling pathways that trigger eIF2α phosphorylation independently of PKR. Examination of rRNA revealed it was largely intact at two hours post-poly(I:C) when translation repression is robust (Figures 6A and S7C). Furthermore, a parallel study to ours demonstrated that cleavage of rRNA by RNase L does not reduce the ability of ribosome to translate in extracts (Rath et al., 2018). These observations argue that rRNA degradation does not drive RNase L-driven translational repression at early times post-dsRNA, which is also supported by our analysis of IFN-β translation (see below).

**Figure 6.**
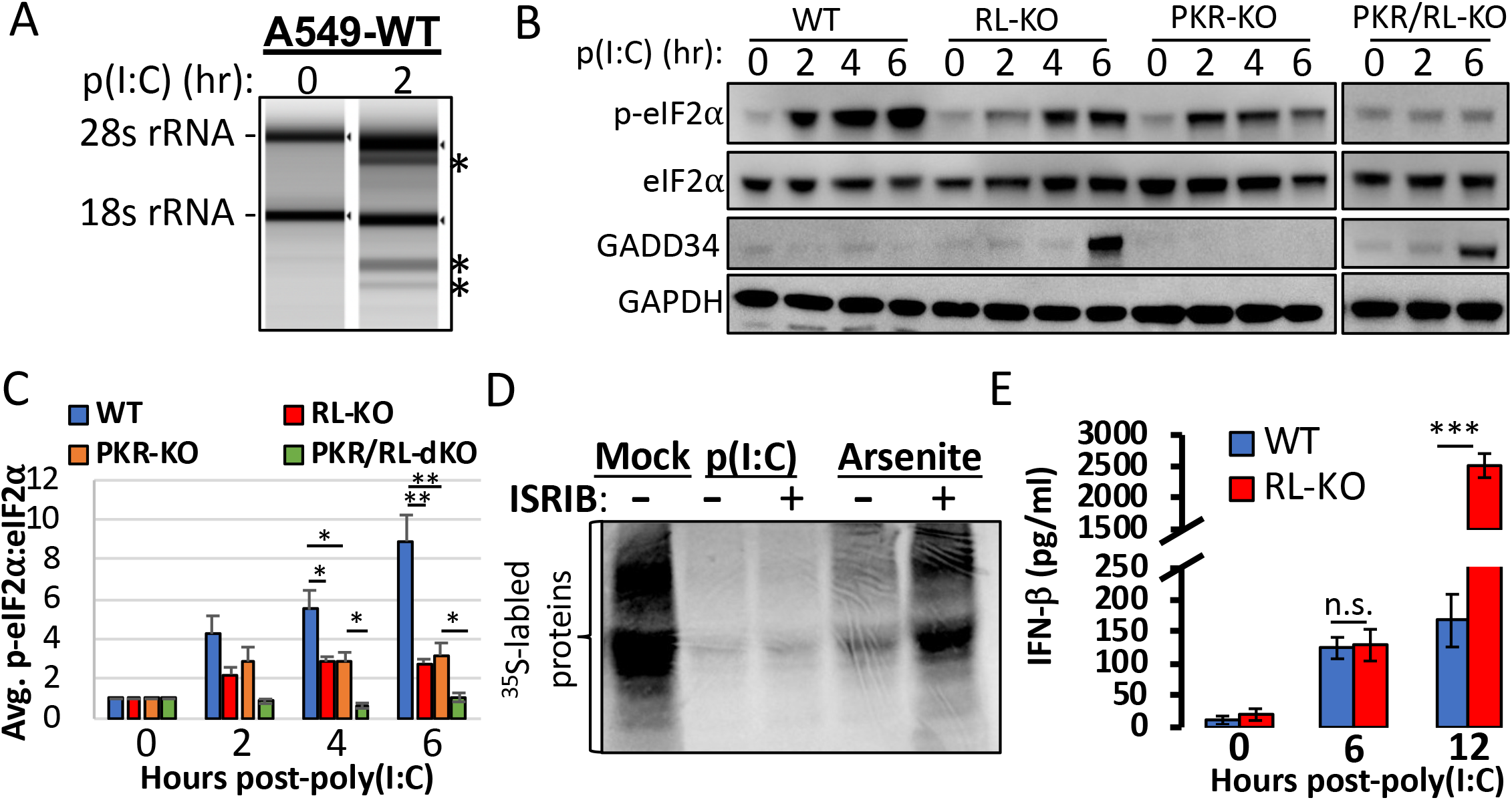
RNase L-driven translational repression is independent of rRNA degradation and p-eIF2α. (A) TapeStation analysis of the 28S and 18S rRNAs. (B) Immunoblot analysis of eIF2α, eIF2α-P51S (p-eIF2α), GADD34, and GAPDH in WT, RL-KO, and PKR-KO, and PKR/RL-dKO cells at the indicated times post-poly(I:C). (C) Quantification of the p-eIF2α:eIF2α ratio as represented in (B). Bars represent the average ratio +/− SEM from independent replicates (n=5-9). (D) S-35 metabolic labeling of newly synthesized proteins in A549 cells transfected with poly(I:C) then subsequently treated with or without 50nM ISRIB. As a positive control, cells were treated with 250 uM sodium arsenite for 30 minutes with or without 50nM ISRIB. (E) Quantification of IFN-β secretion from WT and RL-KO A549 via ELISA. Limit of quantification was 50 pg/ml. Bars represent Average +/− S.D. from three independent experiments. * indicates p-value <0.05, ** indicates p-value < 0.005, and *** indicates p-value <0.001 as determined by student’s t-test.

Examination of eIF2α phosphorylation revealed that RNase L promotes phosphorylation of eIF2α, as p-eIF2α levels were lower (2-3 fold) in cells lacking RNase L activity post-poly(I:C) (Figures 6B,C and S7D,E). This phosphorylation of eIF2α is independent of PKR, since PKR-KO cells also showed eIF2α phosphorylation post-poly(I:C), albeit less than in WT cells (Figures 6B,C and S7F). Furthermore, this alternative mode of eIF2α phosphorylation is dependent on RNase L since eIF2α phosphorylation was abolished to background levels in PKR-KO RL-KO cell lines. This suggests that RNase L activates an additional eIF2α kinase. Moreover, RNase L inhibits the induction of the GADD34 eIF2α phosphatase post-poly(I:C) (Figures 3B, 6B and S6G), which likely promotes elevated p-eIF2α levels. Combined, these data argue that RNase L activation promotes phosphorylation of eIF2α.

To determine whether RNase L-promoted phosphorylation of eIF2α contributes to RNase L-driven translational shut-off, we quantified translational activity in WT cells following transfection of poly(I:C) in the presence or absence of ISRIB, which bypasses the inhibitory effect of p-eIF2α on translation (Sidrauski et al., 2015). Strikingly, poly(I:C)-induced translational arrest was unaffected by ISRIB treatment (Figure 6D). In contrast, ISRIB de-repressed sodium arsenite-induced translation arrest, which occurs through eIF2α phosphorylation. This demonstrates that phosphorylation of eIF2α is not the primary driver of rapid RNase L-mediated translational repression.

### RNase L resistant mRNAs continue to translate during a dsRNA response

The observations above argue RNase L repression of translation is not through rRNA degradation or eIF2α phosphorylation. This suggests that RNase L mediated mRNA turnover accounts for the bulk of translational repression at early times during acute dsRNA stress. Nevertheless, it remained formally possible that translation is repressed by an unknown RNase L-dependent mechanism. We reasoned that if translational repression is simply due to mRNA degradation then mRNAs that escape RNase L degradation, such as IFN-β, should still produce protein. Alternatively, if global translation is repressed by an unknown signaling mechanism or degradation of tRNAs, then IFN-β protein would not be expressed despite the presence of the *IFN-β* mRNA. Given this, we quantified IFN-β secretion from WT and RL-KO cells following poly(I:C) transfection via ELISA.

An important observation is that WT and RL-KO cells excreted equivalent levels of IFN-β at six hours post-poly(I:C) (Figure 6E), indicating that RNase L-driven translational repression at early times post-poly(I:C) does not negatively affect IFN-β translation and secretion. These results demonstrate that RNase L-resistant mRNAs are able to be translated at early times post-dsRNA. This argues that the translation machinery is functional and translation is simply limited by the presence of mRNAs.

Interestingly, by twelve hours post-poly(I:C), secreted IFN-β from WT cells was ~10-fold lower than from RL-KO cells. We suggest this is likely the result of substantial rRNA degradation by RNase L at these later times and/or cells entering the apoptotic pathway (Figure S7C).

## DISCUSSION

Several lines of evidence demonstrate that RNase L leads to a widespread degradation of the majority of cytoplasmic mRNAs. First, we observe by smFISH that RNase L activation leads to dramatic decreases in the cytoplasmic AHNAK and GAPDH mRNAs, as well as the *NORAD* lincRNA (Figures 2A,B,C,D,E and S3G,H). Second, we observe that the *GAPDH, AHNAK, ACTB,* and *TUBA* mRNAs are all reduced in response to poly(I:C) treatment in an RNase L-dependent manner via qRT-PCR (Figures 2C,F). Third, examination of poly(A)+ mRNAs in the cytosol showed an approximately 70% RNase L-dependent reduction in total mRNA levels (Figures 2G and S3J). Finally, RNA-Seq revealed a robust RNase L-dependent reduction in the majority of mRNAs (Figure 4B). Consistent with these findings, a related study demonstrated that RNase L increases decay rates of large numbers of mRNAs in response to dsRNA (Rath et al., 2018).

Despite the promiscuous nature of RNase L degradation, some mRNAs are clearly resistant to RNase L-mediated degradation. For example, the *IFN-β* mRNA is expressed at similar levels in WT and RL-KO cells post-poly(:C) by both qRT-PCR and smFISH, indicating that RNase L does not affect the levels of this mRNA (Figure 3). Indeed, from our RNA-Seq analyses, we identify numerous mRNAs that fully, or partially, escape RNase L-mediated mRNA degradation. Importantly, many of these mRNAs encode for dsRNA-induced antiviral proteins and cytokines (Figure 4C). While the transcriptional induction of these mRNAs may partially contribute to their ability to escape of RNase L-mediated mRNA turnover, several observations suggest that post-transcriptional mechanisms also play a role. First, some IRF3-induced mRNAs (i.e. *IL-6* and *GADD34* mRNAs) do not fully escape RNase L (Figure 3B). Second, abundant levels of IFN-βmRNA are present in the cytosol as early as two hours post-poly(I:C) when RNase L is actively promoting the rapid degradation of mRNAs (Figure S5A). Third, RNase L promotes continuous rRNA degradation up to eight hours post-poly(I:C) (Figure S7C), during which time we observed abundant cytosolic *IFN-β* mRNAs at comparable levels in WT and RL-KO cells (Figure 3A).

Finally, some constitutively expressed mRNAs (i.e. *STAT1 and MYC)* appear to be resistant to RNase L-driven mRNA turnover (Figure 4B). Notably, poliovirus mRNA is resistant to RNase L-mediated cleavage via a conserved RNA structure (Han et al., 2007; Townsend et al., 2008). Therefore, specific host mRNAs may contain structures, elements, or modifications that impart resistance to RNase L-mediated mRNA turnover via yet-to-be-determined mechanisms.

Several of our observations argue that one consequence of RNase L-mediated mRNA degradation is host shut-off of translation. First, we observed that RNase L catalytic activity represses translation by two hours post-dsRNA (Figures 5A,B and S7A,B), when rRNA is largely intact (Figures 6A and S7C), but bulk mRNA levels are substantially reduced (Figures 2 and 4). Second, RNase L-dependent bulk translational repression is unaffected by ISRIB or knockout of PKR (Figures 5A,B, 6D and S7A.B), indicating RNase L-driven translational repression occurs independently of eIF2α phosphorylation. Finally, because the *IFN-β* mRNA escapes RNase L-driven mRNA turnover and IFN-β protein is produced at similar levels in WT and RL-KO cells prior to robust rRNA degradation (Figures 3 and 6E), this demonstrates that the presence of a stable mRNA is sufficient to produce protein, and therefore the translation machinery is functional. Consistent with this latter finding, a related study demonstrated that antiviral cytokines are produced during RNase L-mediated translational arrest (Chitrakar et al., 2018). Combined, these observations reveal a new mode of RNase L-driven translational repression that occurs via bulk mRNA turnover and permits the translation of key RNase L-resistant antiviral mRNAs, such as the *IFN-β* mRNA (Figure 7).

**Figure 7.**
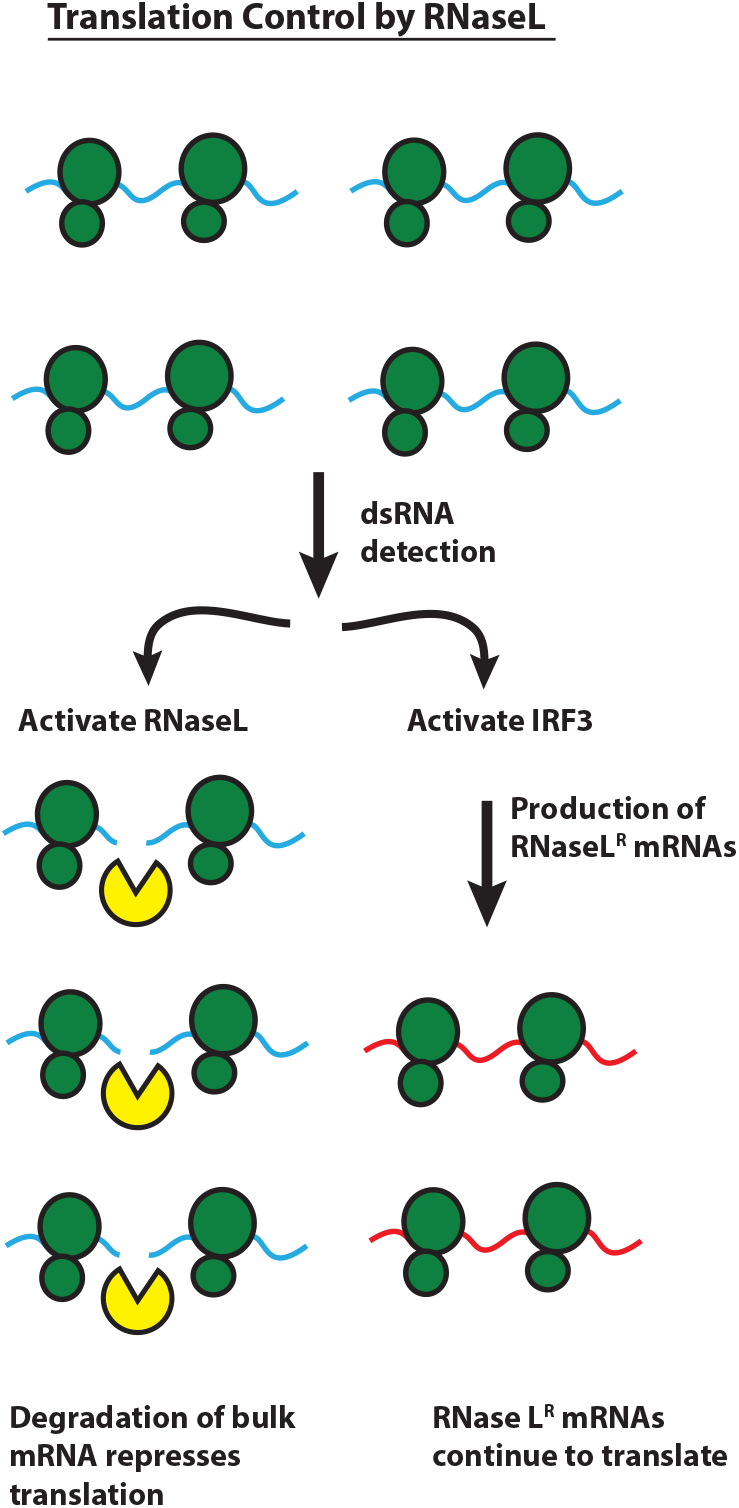
Model of RNase L reprogramming of translation via differential mRNA turnover.

A second consequence of robust mRNA degradation by RNase L is PABPC1 translocation to the nucleus (Figure 1), which is known to occur during global mRNA degradation and viral replication (Glaunsinger and Ganem, 2004, Kumar et al., 2011, Gray et al., 2015, Montero et al., 2008, Dobrikova et al., 2010, Borah et al., 2011). Since PABPC1 is core component of SGs, this may contribute to the RNase L-mediated regulation of SG assembly/composition. More importantly, translocation of PABPC1 to the nucleus can inhibit global transcription of host genes (Gilbertson et al., 2018), though this may not apply to IRF3-induced genes since their transcription increases in the presence of nuclear PABPC1 accumulation (Figures 1,3, and 4). These observations suggest a possible new means by which RNase L contributes to host shutoff, whereby RNase L-promoted PABPC1 translocation alters transcription.

An additional consequence of RNase L-mediated mRNA turnover is the inhibition of canonical PKR-dependent SG assembly in response to dsRNA (Figures 1, 2 and S3G). These observations reinforce recent findings that RNA is required for SG assembly (Van Treek et al. 2018). Large canonical SGs are often observed in response to dsRNA or viral replication (Onomoto et al., 2012). We suggest that this is due to the cells being assayed, such as MEFs, have a weak RNase L response (Li et al., 2017, Banerjee et al., 2014), the virus being assayed inhibiting the OAS/RNase L pathway, and/or the kinetics and/or dose of dsRNA production during viral replication may promote PKR-mediated SG assembly prior to RNase L activation. Nevertheless, small punctate SG morphologies and PABPC1 translocation, hallmarks of RNase L activation revealed in our studies, have been observed during Rotavirus replication, which triggers the OAS/RNase L pathway (Montero et al., 2008). This suggests that RNase L may promote RLB formation and prevent SG formation during some viral infections. Since SGs are proposed to promote antiviral signaling and are pro-survival, and RLBs contain numerous proteins involved in mRNA metabolism and processing (Figure S2C), an important issue in future work will be to determine the functions RNase L-mediated inhibition of SGs and the formation of RLBs during the dsRNA/antiviral response.

We suggest that the activation of RNase L and widespread degradation of host mRNAs acts in concert with other aspects of the dsRNA response. Specifically, the sensing of dsRNA by PRRs leads to: 1) the activation of IRF3, which promotes the transcription of antiviral genes such as type I interferons; 2) the activation of RNase L, which initiates widespread degradation of both host and viral RNAs; 3) PKR-and RNase L-mediated promotion of eIF2α phosphorylation changing the specificity of translation initiation; 4) Inhibition of canonical translation re-initiation via RNase L-mediated downregulation of the GADD34 p-eIF2α phosphatase (Figures 3B, 6B, and S7G). In the context of these four outcomes, the majority of IRF3 induced genes appear to be resistant to RNase L-mediated mRNA turnover, and this allows for their selective translation. Thus, RNase L-mediated degradation of cellular mRNAs leads to reprograming of the produced proteins to only those mRNAs resistant to RNase L activity.

Strikingly, by examining single cells, we observe that different aspects of the cellular response to dsRNA can be triggered independently in individual cells. We observe numerous instances where RNase L activation and PKR activation are uncoupled from induction of IRF3 targets, such as the *IFN-β* mRNA (Figures 3A and S4A,C). The ability to independently activate different arms of the cellular response to dsRNA allows cells to modulate their transcriptional and translational output, as well as to determine whether to enter apoptosis. An important aspect of future work will be to determine the additional inputs that affect the independent activation of these pathways and how those affect the cellular response to viral infection

## Supporting information

## ACKNOWLEDGMENTS

The authors would like to thank Dr. Christopher Sullivan for the pLenti-CMV-RNase L, pLenti-RNase L-R667A, and pLenti-EF1-blast vectors and the A549 cell line. We thank Dr. Nancy Kedersha for the U-2 OS cell line. We thank Dr. Susan Weiss for the A549-PKR-KO cell line. This work was supported by funds from HHMI.

## AUTHOR CONTRIBUTIONS

JMB and RP conceived the project. JMB, SLM, ETL, and TM performed experiments. JMB, SLM, ETL, TM and RP reviewed and interpreted the data. JMB and RP wrote the manuscript.

## STAR METHODS

### Plasmids

To generate the 3xflag-tagged RNase L expression vectors used for transient transfections, the RNase L and RNase L-R667 coding sequences were tagged with 3x-flag encoding sequence via PCR amplification from the pLenti-CMV-RNase L and pLenti-RNase L-R667A vectors using the RL-3xFlag-Kpn1/Xho1 primers (sense: TCGAGGTACCATGGAGAGCAGGGATCATAA; antisense:

AGCTCTCGAGTCAGCACCCAGGGCTGGCCA)primers and Phusion polymerase (New England BioLabs). The PCR amplicons were digested with Kpn1 and Xho1 and ligated into the *Kpn1/Xho1* sites of pcDNA3.1-puro using T4 DNA ligase (New England BioLabs). To generate pLenti-EF1-Blast lentiviral genomes encoding RNase L and RNase L-R667A, the RNase L and RNase L-R667A coding sequences were N-terminally-tagged with 3x-flag tag via PCR amplification expression vectors using RL-3x-flag-fusion primers: (RL_sen_3x_FLAG: CATGGACTACAAAGACCATGACGGTGATTATAAAGATCATGACATCGACTACAAGGATGACGATGACAAGATGGAGAGCAGGGATCATAA; RL_sen_kpn 1 _kozak_flag: GTGGGTACCGCCACCATGGACTACAAAGACCATGACGGTG; RL_anti_EcoN1: GAAGAACCTTCACAAGGGAATGGTCATAAT) and inserted into the *Xho1/EcoN1* sites of the pLenti-EF1-Blast vector using In-Fusion (Clontech). To generate the pLJM1-GFP-G3BP1 lentiviral vector, the GPF-G3BP1 coding sequence was subcloned from the pEGFP-C1-G3BP1+stop vector, a gift from Dr. Nancy Kedersha, via digestion with Nhe1 and EcoR1 and ligation into the pLJM1-EGFP lentiviral vector (Addgene: 19319) using T4 DNA ligase (New England BioLabs). The Lentiviral packaging vectors – pVSV-G, pRSV-Rev, and pMDLg/pRRE – were a gift from Dr. Sabrina Spencer. The px458 Cas9 vector (Addgene: 48138) was used to generate knockout cell lines. The CRISPR/Cas9 guide RNAs were designed using the Integrated DNA Technologies (IDT) CRISPR guide target design tool. Overlapping oligos (RL sgRNA 1 sense: CACCGCGCATCTGCTGCTGGACCA; RL sgRNA 1 antisense: AAACTGGTCCAGCAGCAGATGCGC) were annealed in T4 DNA ligase buffer and ligated into the *Bbs1* sites in px458 using T4 DNA ligase. All plasmids were sequence verified via sanger sequencing (Quintarabio).

### Antibodies

Mouse anti-RNase L Antibody 2E9 (Novus Biologicals: NB100-351) was use at 1:1500 for immunoblot analyses. Mouse monoclonal anti-G3BP antibody (Abcam: ab56574) was used at 1:500 for IFA and 1:1000 for IB analyses. Rabbit anti-EIF2S1 (Phospho S51-eIF2α) monoclonal antibody (Abcam: ab32157) was used at 1:500 for IB analysis. Rabbit anti-eIF2α (Cell Signaling Technology: CST9722S) was used at 1:1000 for IB analysis. Mouse monoclonal anti-FLAG (Sigma Aldrich: F1804) was used at 1:1000 for IB analysis. Rabbit anti-GAPDH (Cell Signaling Technology: 2118L) was used at 1:2000 for IB analysis. Rabbit anti-PKR (Cell Signaling Technology: 12297S) was used at 1:1000 for IB analysis. Rabbit polyclonal anti-TIA1 (Abcam: ab40693) was used at 1:500 for IFA. Rabbit anti-FAM120A (Sigma-Aldrich: HPA019734) was used at 1:500 for IFA. Rabbit polyclonal anti-Caprin1 (Fisher Scientific: 50554-357) was used at 1:500 for IFA. Rabbit polyclonal anti-PABP antibody (Abcam: ab21060) was used at 1:500 for IFA. DCP1B (D2P9W) antibody (Cell Signaling: 13233) was used at 1:500 for IFA as a p-body marker. Goat anti-mouse IgG H&L FITC (Abcam: ab97022) was used at 1:1000 for IFA. Goat antirabbit IgG H&L Alexa Fluor 647 (ab150079) was used at 1:1000 for IFA. Anti-rabbit IgG, HRP-linked antibody (Cell Signaling Technology: 7074S) was used at 1:3000 for IB analysis. Anti-mouse IgG, HRP-linked antibody (Cell Signaling Technology: 7076S) was used at 1:10,000 for IB analysis. Anti-puromycin (Millipor Sigma: MABE343) was used at 1:1000 for SUnSET analyses.

### Cell culture

Cells were maintained at 5% CO2 and 37 degrees Celsius in Dulbecco’s modified eagle’ medium (DMEM) supplemented with fetal bovine serum (FBS; 10% v/v) and penicillin/streptomycin (1% v/v). Cells were routinely tested for mycoplasma contamination by the cell culture core facility and were negative for mycoplasma contamination throughout the study. Cells were transfected with poly(I:C) HMW (InvivoGen: tlrl-pic) using 3-ul of lipofectamine 2000 (Thermo Fisher Scientific) per 1-ug or poly(I:C).

### Generation of knockout cell lines

To generate RNase L knockout A549 and U-2 OS lines, cells (T-25 flask; 70% confluent) were cotransfected with 2-ug of px458-RL and 200-ng of pcDNA3.1-puro using 6-ul of Lipofectamine 2000 (Thermo Fisher Scientific) according to manufacturer’s instructions. Twenty-four hours post-transfection after Cas9-GFP expression was observed via fluorescent microscopy, the medium was replaced with medium containing 2ug/ml of puromycin. Selective medium was replaced 3 days post-transfection. Five days post-transfection, selective growth medium was replaced with normal growth medium. When cells became 80% confluent, cells were serial diluted and plated on 15-cm dishes. Individual colonies were isolated, propagated, and screened via immunoblot analysis.

### Generation of Lentiviral particles

To generate GFP-G3BP1 lentiviral particles, HEK293T cells (15-cm dish; 80% confluent) were co-transfected with 11.7-ug of pLMJ1-GFP-G3BP1, 3.5-ug of pVSV-G, 2.9-ug of pRSV-Rev, and 5.6-ug of pMDLg-pRRE using 100-ul of lipofectamine 2000. Medium was replaced 6 hours post-transfection. Medium was collected at twenty-four and forty-eight hours post-transfection and filter-sterilized with a 0.45-um filter. The RNase L and RNase L-R667A lentiviral particles were generated via the same methods using the pLenti-EF1-RNase L-blast and pLenti-EF1-3xflag-RNase L-R667A-blast.

### Generation of stable cell lines

To reconstitute A549-RL-KO cells with either 3xflag-RNase L or 3xflag-RNase L-R667A, A549-RL-KO cells were seeded in T-25 flask. When 80% confluent, cells were incubated for 1 hour with 1-ml of either 3xflag-RNase L or 3xflag-RNase L-R667A lentiviral particles containing 10-ug of polybrene with periodic rocking. Normal medium was then added to the flask and incubated for twenty-four hours. Medium was removed 24 hours post-transduction and replaced with selective growth medium containing 10-ug/ml of Blasticidine S hydrochloride (Sigma-Aldrich). Selective medium was changed every three days. After one-week, selective medium was replaced with normal growth medium. Expression of RNase L was confirmed via immunoblot analysis. To generate WT and RL-KO U-2 OS cells that constitutively express GFP-G3BP1, cells (T-25 flask, 80% confluent) were incubated with 1-ml GFP-G3BP1 lentivirus particles (6.4 x 10^5^ IU/ml; MOI~0.5) containing 10-ug/ml of polybrene for one hour. Normal medium was then added to the flask. 24 hours post-transduction, cells were re-seeded in T-25 flask containing 2-ug/ml Puromycin selective medium. Cells were maintained in selective medium for four days before returning to normal growth medium. GFP-G3BP1 expression was confirmed via fluorescent microscopy and IB analysis.

### Immunoblot analyses

To screen and confirm for knockout or reconstitution of proteins, cells were lysed in SDS solution (1% SDS, 2% β-mercaptoethanol) by boiling for 10 min followed by 1 min of vortexing. Equal volumes of lysates were fractionated on 4-12% Bis-Tris Protein gels (Thermo Fisher Scientific) in MOPS buffer and transferred to nitrocellulose membrane (GE Healthcare). Membranes were blocked in 5% BSA in TBST. Membranes were then incubated with primary antibodies for 2 hours at room temperature or overnight at 4 degrees Celsius. After washing, membranes were incubated with HRP-linked anti-rabbit IgG or anti-mouse IgG secondary antibodies for 1 hour at room temperature. After washing, membranes were incubated with ECL substrates (Thermo Fisher Scientific: 32106) for 1-5 minutes. Membranes were then stripped using Restore western blot stripping buffer (Thermo Fisher Scientific: 21059) and re-blocked with 5% BSA in TBST. Membranes were then incubated with anti-GAPDH, washed, incubated with HRP-linked anti-mouse antibody, washed, and incubated in ECL substrate. For quantitation of p-eIF2α, cells were seeded in 12-well format. Cells were transfected with 250-ng of poly(I:C) and cell lysates were collected at indicated time post-transfection. A fifth of the lysate (12-ul/60-ul) was fractionated on 4-12% Bis-Tris Protein gels in MOPS buffer and transferred to nitrocellulose membrane. Membranes were blocked in 5% MILK in TBST. Membranes were then incubated with rabbit anti-EIF2S 1 (Phospho S51-eIF2α) antibodies overnight at 4 degrees Celsius. After washing, membranes were incubated with HRP-linked anti-rabbit antibody for 1 hour at room temperature. After washing, membranes were incubated with SuperSignal West Femto Maximum Sensitivity ECL substrate (Thermo Fisher Scientific: 34095) for 1-5 minutes. Membranes were stripped and re-probed using with rabbit anti-eIF2α, washed, incubated with HRP-linked anti-rabbit antibody for 1 hour at room temperature, washed, incubated with SuperSignal West Femto Maximum Sensitivity ECL substrate for 1-5 minutes. Photographs of membranes were taken using ImageQuant LAS 4000 (GE Healthcare) and analyzed using ImageJ with Fiji plug-in. For quantitation of western blots, average band signal was determined using the measure function in imageJ.

### Sequential immunofluorescence and single molecule FISH

Sequential immunofluorescence and smFISH was performed following manufacturer’s protocol (https://biosearchassets.blob.core.windows.net/assets/bti_custom_stellaris_immunofluorescence_seq_prot_ocol.pdf). Ship ready GAPDH smFISH probes labeled with Quasar 570 Dye were purchased from Stellaris. Custom AHNAK and NORAD smFISH probes labeled with Quasar 670 dye were designed and purchased from Stellaris and are described in (Khong et al., 2017). Custom IFNB1 smFISH probes were designed using Stellaris smFISH probe designer (Biosearch Technologies) available online at http://www.biosearchtech.com/stellaris-designer. Reverse complement DNA oligos were purchased from IDT. The IFNB1 smFISH probes were labeled with Atto-633 using ddUTP-Atto633 (Axxora: JBS-NU-1619-633) and terminal deoxynucleotidyl transferase (Thermo Fisher Scientific: EP0161) as described in (Gaspar et al., 2017). Oligo d(T)30-Cy3 were purchased from IDT.

### Microscopy and Image Analysis

Immunofluorescence and smFISH with DAPI staining were imaged using a wide field DeltaVision Elite microscope with a 100X objective using a PCO Edge sCMOS camera. For IFA, 10 Z sections at 0.3 um/section were taken for each image. For IFA/smFISH, 15 Z planes at 0.2 um/section were taken for each image. Deconvoluted images were processed using ImageJ with FIJI plugin. Z-planes were stacked and minimum and maximum display values were set in ImageJ for each channel to properly view fluorescence. Quantification SGs was determined using Imaris Image Analysis Software (Bitplane) (University of Colorado-Boulder, BioFrontiers Advanced Light Microscopy Core). Live cell imaging was performed using a Nikon Spinning Disk Confocal microscope outfitted with an environmental chamber with O2, CO2, temperature, and humidity control (University of Colorado-Boulder, BioFrontiers Advanced Light Microscopy Core). All images were acquired using a 2x Andor Ultra 888 EMCCD camera.

### Mass spectrometry

U2OS cells were grown to 80% confluence in 15cm dishes (two dishes per replicate). Cells were transfected with poly(I:C) at 0.5ug/ml. Four hours post-transfection, the media was then aspirated, cells were resuspended in media, scraped into a 50mL conical, and pelleted via centrifuged at 1,500g. The supernatant was aspirated and the pellets were snap frozen in liquid nitrogen. After thawing, cells were resuspended in 1mL of stress granule lysis buffer (50mM Tris HCL pH 7.4, 100mM Potassium Acetate, 2mM Magnesium Acetate, 0.5mM DTT, 50ug/ml Heparin, 0.5% NP40, 1 complete mini EDTA free protease inhibitor tablet per 50mL of buffer). Cells were then passed through a 25G 5/8 needle 7 times on ice to lyse. At this step, lysate was inspected by wide field microscopy to determine if granules were visible in the media. Cells were then pelleted by centrifugation at 300g for 5 minutes at 4 deg. The supernatant was taken RNP complexes were pelleted via centrifugation at 18,000g for 20 minutes at 4 deg. The pellet was resuspended in 1mL of stress granule lysis buffer and pelleted via centrifugation at 18,000g for 20 minutes at 4 deg. To preclear the samples and remove non-specific binders, the pellet was resuspended in 340uL of lysis buffer and 60uL of pre-washed Protein A Dynabeads were added and incubated for 30 minutes at 4 deg on nutator. Dynabeads were than taken off 2x using a magnet and the preclearance step was repeated. Following final removal of beads, 1ug of either anti-GFP antibody (Invitrogen A11122) or anti IgG (Invitrogen 10500C) were added to the respective samples and incubated overnight on nutator at 4 degrees.

Following incubation, samples were centrifuged at 18,000g for 20 min at 4 deg to remove antibody. The pellet was resuspended in 500uL of stress granule lysis buffer and 33uL of washed Protein A Dynabeads (1mg) were added and nutated for 3 hours at 4 degrees. The beads were then washed for 2 minutes in wash buffer 1 (stress granule lysis buffer + 2M Urea), 5 minutes in wash buffer 2 at 4 degrees (stress granule lysis buffer + 300mM of potassium acetate), 5 minutes with stress granule lysis buffer at 4 degrees. The sample was then washed 8x with 1mL of TE buffer to remove detergent and beads were brought up in 50uL of TE buffer.

Samples were then processed by the mass spectrometry facility at CU Boulder and analyzed on Thermo LTQ Orbitrap (Thermo Fisher Scientific). Proteins with fewer then 5 cumulative spectral counts between the three replicates were removed. The spectral counts from the remaining proteins were averaged and divided by the spectral counts in the IgG control. Proteins that were two-fold enriched over the IgG control were selected for further analysis. A stress granule reference file was created by merging the proteins identified in three different stress granule proteomic studies that stress cells with sodium arsenite (Jain et al., 2016, Youn et al., 2018, Mol Cell, Markmiller et al., 2018), which resulted in a stress granule proteome of 491 proteins. To determine the overlap between the poly I:C granule proteome and the sodium arsenite stress granule proteome, the two protein lists were inner joined using R. Gene Ontology was performed on the proteins that did not overlap with the stress granule proteome. Gene ontology biological processes were derived from Gene Ontology Consortium enrichment analysis (Mi et al., 2016; http://www.geneontology.org/).

### RT-qPCR

WT and RL-KO A549 cells (12-well; 60% confluent) were transfected with or without poly(I:C). Six hours post-transfection, RNA was extracted, treated with DNase I (NEB) for 15 minutes, and re-purified via ethanol (75%) sodium acetate (0.3M) precipitation, and re-suspended in 15-ul of water. Equal volumes (400-ng of WT RNA) were then reverse transcribed using super script III reverse transcriptase (Thermo Fisher Scientific) and polydT(20) primer (Integrated DNA Technologies). cDNA was diluted to 100-ul. 2-ul of cDNA was added to qPCR reaction containing iQ SYBR green master mix (Bio-Rad) and 10-pmol of gene-specific primers (GAPDH sense: CTTTGGTATCGTGGAAGGACTC; GAPDH antisense: GTAGAGGCAGGGATGATGTTC; IFN-B sense: GCCGCATTGACCATCTATGA, IFN-B antisense: GCCAGGAGGTTCTCAACAATAG; ACTB sense: CAGGAAGTCCCTTGCCATCC; ACTB antisense: TGCTATCACCTCCCCTGTGT; TUBA1A sense: GTGAGTTTTCAGAGGCCCGT; TUBA1A antisense: AAAGCAGCACCTTTGTGACG; IL6_sen: ATCTAGATGCAATAACCACCCCT; IL6_anti: AGCTGCGCAGAATGAGATGA; GADD34_sense: GAAACCCCTACTCATGATCCG; GADD34_anti: AAATGGACAGTGACCTTCTCG). Reactions were run in triplicate on CFX96 qPCR machine (Bio-Rad) using standard two-step cycle. PCR fragment sizes were confirmed by ethidium bromide staining and RT-controls were included to demonstrate that amplification was from cDNA and not gDNA.

### Metabolic labeling of newly synthesized proteins

Wild-type, RNase L knockout or PKR knockout A549 cells were transfected with poly(I:C) as described above. S35 metabolic labeling of nascent proteins was performed as described in Moon and Parker, 2018. Briefly, cells were incubated with ^35^S labeled met and cys (EXPRE35S35S Protein Labeling Mix, PerkinElmer) in labeling medium (DMEM lacking met and cys (Sigma Aldrich) supplemented with 10% dialyzed FBS (Sigma Aldrich), glutaMAX (Gibco) and 1% streptomycin/penicillin) for 30 minutes at two-and four-hours post-transfection following a 30 minute incubation in labeling medium to deplete intracellular amino acid stores. Cells were harvested in NP-40 lysis buffer with protease and phosphatase inhibitors (Cell Signaling Technologies), lysed and equal volumes of lysate run on NuPAGE 4-12% Bis-Tris protein gels (Thermo Fisher Scientific). Gels were exposed to phosphor screens and imaged on a typhoon FLA 9500 phosphorimager. The average relative translation activity was determined using ImageJ (Fiji) (Schindelin et al., 2012) to quantify total lane intensity for each sample from 2-3 independent experiments.

SUnSET was performed as described by Schmidt et al., 2009. Briefly, puromycin (10 ug/ml) was added to cells 10 minutes prior to fixing or harvesting lysates. For IF of puro-labeled cells, cells were imaged on 60X objective. Between 5 and 10 fields of view were imaged. The fluorescent intensity of puromycin labeling of individual cells was measured using fiji software.

### IFN-β ELISA

WT and RL-KO A549 cells (6-well format, 1 ml or medium, 70% confluency) were transfected with poly(I:C). At six- and twelve-hours post-poly(I:C), 50-ul of medium was removed from well and immediately assayed via ELISA using IFN beta human ELISA kit (Thermo Fisher Scientific; 414101) using manufacturer’s instructions. Time zero was taken by removing 50-ul of medium prior to poly(I:C) transfection.

### High-throughput RNA sequencing

WT and RL-KO A549 cells (100-mm dish, 60% confluent) were transfected with or without 500-ng/ml of poly(I:C). Total RNA was isolated six hours post-transfection. The ERCC spike-in was added according to manufacturer’s instructions and RNA libraries were prepared by the BioFrontiers Sequencing Core at CU Boulder. Libraries were sequenced on Illumina NextSeq 500. Read quality was assessed using fastqc. Illumina adaptors were trimmed using Trimmomatic 0.32 in paired and (PE) mode (Bolger et al., 2014). An index genome was acquired from GENCODE (Release 19 GRCh37.p13). Reads were aligned using Tophat (version 2.0.6) and Bowtie2 (version 2.0.2) and the following parameters:-b2 -fast – microexon-search (Kim et al., 2013; Langmead and Salzberg, 2012). Differential expression analysis was performed using Cuffdiff (version 2.2.1) with the default parameters (Trapnell et al., 2013). Gene ontology biological processes were derived from Gene Ontology Consortium enrichment analysis (Mi et al., 2016; *http://www.geneontology.org/*). MEME analyses were performed using MEME suite (Bailey et al, 2009). Meta-analysis of CLIP sites were derived from Yang et al., 2015. Ensembl Biomart was used for retrieval of transcript characterization (Kinsella et al., 2011). Au-rich elements were predicted using ARED (https://brp.kfshrc.edu.sa/ared/Home/BasicSearch) (Bakheet et al., 2018). The RNA-seq data files have been submitted to GEO database (Accession ID will be added).

## SUPPLEMENTAL FIGURE LEGENDS

**Supplemental Figure S1** (related to Figure 1). (A) Western blot analysis confirming knockout of RNase L expression in A549 and U-2 OS clonal cell lines used for analysis in this study. (B) Western blot showing rescue of RNase L expression in the A549-RL-KO cell line via lentiviral transduction. Note that the reconstituted RNase L migrates slightly slower than the endogenous protein due to the N-terminal 3x-flag tag, and that expression is ~ten-fold higher than the endogenous protein. (C) Western blot showing rescue of RNase L expression in the A549-RL-KO cell line via transient transfection of expression vectors encoding untagged RNase L. (D) Images of individual staining of G3BP1 and PABPC1 via IFA for the merged image shown in Figure 1C. (E) IF for G3BP1 (green) and PABPC1 (red) in A549-RL-KO cells transfected with pcDNA3.1 empty vector (pEV), pcDNA3.1-RNase L (pRL), or pcDNA3.1-RNase L-R667A (pRL-CM) twenty-four hours prior to transfection with poly(I:C). (F) IF G3BP1 (green) and PABPC1 (red) in WT and RL-KO A549 cells one hour post-sodium arsenite treatment (500 uM).

**Supplemental Figure S2** (related to Figure 1). (A) IF for indicated SG-associated proteins in WT A549 cells six hours post-transfection of poly(I:C). (B) IF for indicated SG-associated proteins in RL-KO A549 cells six hours post-transfection of poly(I:C). Scale bars represent 15 um. (C) Gene ontology of RLB-associated proteins identified by mass spectrometry.

**Supplemental Figure S3** (related to Figures 1&2). (A) FISH for poly(A)+ RNA in WT cells transfected with poly (I:C). (B) Live cell imaging of U-2 OS-GPF-G3BP1 cells. Cells were treated with 500 uM sodium arsenite followed by treatment with or without 50ug/ml cycloheximide, and images were taken ninety minutes post-treatment. For poly(I:C) treatment, cells were transfected with 500ng/ml of poly (I:C). Forty-five minutes later before SGs had formed, cells were treated with cycloheximide. Images were taken ninety minutes post-transfection of poly(I:C). Scale bar represents 10 um. (C) Western blot analysis of PKR in WT, PKR-KO, and RL-KO cells. (D) Western blot to confirm knockout of RNase L in the A549 PKR-KO cell line. Clones 14 and 17 do not express RNase L. (E) Western blot to confirm rescue of stable 3xflag-RNase L expression in PKR/RL double knockout cells via lentiviral transduction. (F) Individual images for G3BP1 and PABPC1 IFA shown in merged image in Figure 1F. (G) smFISH/IF for NORAD lcnRNA (red) and G3BP1 (green) in WT and RL-KO U-2 OS cells transfected with or without (mock) 500-ng/ml of poly(I:C). (H) Quantification of NORAD smFISH foci as represented by the images on the left. Greater than 14 cells from three different fields were used for analysis. (I) Quantification of the change in *NORAD* levels via RT-qPCR analysis in WT and RL-KO A549 cells six-hours post-poly(I:C) transfection. (J) Quantification of poly(dT) staining in WT and RL-KO cells that contain poly(I:C) induced SGs at the indicated time post-poly(I:C) transfection. Mean intensity was normalized to mock-treated cells. The bars represent mean intensity +/− S.D. from greater than 20 cells from at least 3 separate fields of view.

**Supplemental Figure S4** (related to Figures 3). (A) smFISH/IF for *IFN-β* mRNA (green), *GAPDH* mRNA (red), and G3BP1 (cyan) in WT A549 cells two hours post-transfection of poly(I:C). (B) smFISH/IF for *IL-6* mRNA (green), *GAPDH* mRNA (red), and G3BP1 (cyan) in WT and RL-KO A549 cells six hours post-transfection of poly(I:C). (C) smFISH for *IFN-β* mRNA (red) and IF for G3BP1 (cyan) in WT A549 cells six-hours post-transfection of poly(I:C). Individual cells are demarcated by lines. The presence (+) and absence (-) of RLBs and *IFN-β* mRNA are indicated in each cell. The field shown contains cells in which all combinations of *IFN-β* mRNA and RLB presence/absence were observed to illustrate the heterogeneity of the dsRNA response at the individual cell level and to demonstrate that SG assembly is not necessarily coupled to *IFN-β* induction and vice versa. Scale bars represent 15 um.

**Supplemental Figure S5** (related to Figure 4) (A) Scatterplot showing RNase L-KO FPKM vs. WT FPKM in untreated conditions. Red dots indicate differentially expressed genes (Fold change > 2 or Fold change < .5, p-value < .05). Gray dots indicate ERCC spike-in control. Dark gray trendline indicates correlation for ERCC spike-in control. Light gray trendline indicates correlation for all transcripts. (B) Scatterplot depicting poly(I:C) FPKM values vs. untreated FPKM values in RNase L knockout cells. Dark gray trendline indicates ERCC spike-in control. Light gray trendline indicates correlation for all transcripts. (C) Same as B, but for wild type cells. (D) Same as B, but color-coded for fold change values based on ERCC spike-in normalization. (E) Same as C, but color-coded for fold change values based on ERCC spike-in normalization.

**Supplemental Figure S6** (related to Figure 4)

(A) Normalized log2(poly(I:C)/untreated) vs. expression levels (FPKM) in wild type cells. (B) Same as A, but for RNase L knockout cells. (C) Normalized log2(poly(I:C)/untreated) in wild type cells vs. RNA half-life. Half life data was obtained from Tani et al. 2012. (D) Same as C, but for RNase L knockout cells. (E) Scatterplot depicting half life vs. expression levels (FPKM). Each transcript is color-coded by its log2(poly(I:C)/untreated) values in wild type cells.

**Supplemental Figure S7** (related to Figures 5 and 6). (A) Western Blot for puromycin pulse-labeled proteins in WT, RL-KO, and PKR-KO cells two-hours post-poly(I:C) (500ng/ml) transfection. (B) Quantification of western blot represented in (A). Bars represent the mean puromycin intensity +/− S.E.M. from four independent replicates (n=4) normalized to GAPDH band intensity and mock treatment. (C) TapeStation analysis of rRNA in WT and RL-KO A549 cells at indicated times post-transfection of poly(I:C). (D and E) Western blot of p-eIF2α and eIF2α in A549 RL-KO cells stably reconstituted with RNase L (A549-RL-KO-RL) or RNase L-R667A (A549-RL-KO-RL-CM) at the indicated times posttransfection of poly(I:C). (F) Western blot of p-eIF2α and eIF2α in WT, PKR, and PKR/RL double KO clones 14 and 17 at the indicated times post-transfection of poly(I:C). the p-eIF2α:eIF2α ratio is shown below. (G) Quantification of GADD34 levels in WT and RL-KO cells at the indicated times post-poly(I:C). Bars represent mean +/− S.D. from four independent experiments.

## REFERENCES

Andersen, J.B., Mazan-Mamczarz, K., Zhan, M., Gorospe, M., and Hassel, B.A. (2009). Ribosomal protein mRNAs are primary targets of regulation in RNase-L-induced senescence. RNA Biol. 6, 305–315.

Bailey, T. L., Boden, M., Buske, F.A., Frith, M., Grant, C.E., Clementi, L., Ren, J., Li, W.W., and Noble, W.S. (2009). MEME SUITE: tools for motif discovery and searching. Nucleic acids res. 37, W202–8.

Bakheet, T., Hitti, E., and Khabar, K.S.A. (2018). ARED-Plus: an updated and expanded database of AU-rich element-containing mRNAs and pre-mRNAs. Nucleic Acids Res. 46, D218–D220.

Banerjee, S., Chakrabarti, A., Jha, B.K., Weiss, S.R., and Silverman, R.H. (2014). Cell-Type-Specific Effects of RNase L on Viral Induction of Beta Interferon. mBio. 5, :e00856–14.

Bolger, A.M., Lohse, M., and Usadel, B. (2014). Trimmomatic: a flexible trimmer for Illumina sequence data. Bioinformatics (Oxford, England) 30, 2114–20.

Borah, S., Darricarrère, N., Darnell, A., Myoung, J., and Steitz, J.A. (2011). A Viral Nuclear Noncoding RNA Binds Re-localized Poly(A) Binding Protein and Is Required for Late KSHV Gene Expression. PLoS Pathog. 7, e1002300.

Brennan-Laun, S.E., Ezelle, H.J., Li, X-L., and Hassel, B.A. (2014). RNase-L Control of Cellular mRNAs: Roles in Biologic Functions and Mechanisms of Substrate Targeting. . J. Interf. Cytoki. Res. 34, 275–288.

Buchan, J.R., and Parker, R. (2009). Eukaryotic Stress Granules: The Ins and Out of Translation. Mol.Cell 36, 932.

Chakrabarti, A., Jha, B.K., and Silverman, R.H. (2011). New Insights into the Role of RNase L in Innate Immunity. J. Interf. Cytoki. Res. 31, 49–57.

Chitrakar, A., Rath, S., Donovan, J., Demarest, K., Li, Y., Sridhar, R.R., Weiss, S., Kotenko, S., Wingreen, N., and Korennykh, A. (2018). Realtime 2-5A kinetics suggests interferons β and λ evade global arrest of translation by RNase L. BioRxiv 476341, doi: https://doi.org/10.1101/47634.

Clemens, M.J., and Williams, B.R. 1978. Inhibition of cell-free protein synthesis by pppA2’p5’A2’p5’A: a novel oligonucleotide synthesized by interferon-treated L cell extracts. Cell 13, 565–572.

Dalet, A., Argüello, R. J., Combes, A., Spinelli, L., Jaeger, S., Fallet, M., Vu Manh, T.P., Mendes, A., Perego, J., Reverendo, M. et al. (2017). Protein synthesis inhibition and GADD34 control IFN-β heterogeneous expression in response to dsRNA. EMBO J. 36, 761–782.

Dalet, A., Gatti, E., and Pierre, P. (2015). Integration of PKR-dependent translation inhibition with innate immunity is required for a coordinated anti-viral response. FEBS Lett. 589, 1539–45.

Dobrikova, E., Shveygert, M., Walters, R., and Gromeier, M. (2010). Herpes Simplex Virus Proteins ICP27 and UL47 Associate with Polyadenylate-Binding Protein and Control Its Subcellular Distribution. J. Virol. 84, 270–279.

Donovan, J., Rath, S., Kolet-Mandrikov, D., and Korennykh, A. (2017). Rapid RNase L-Driven Arrest of Protein Synthesis in the dsRNA Response without Degradation of Translation Machinery. RNA 23, 1660–1671

García, M. A., Gil, J., Ventoso, I., Guerra, S., Domingo, E., Rivas, C., and Esteban, M. (2006). Impact of Protein Kinase PKR in Cell Biology: from Antiviral to Antiproliferative Action. Micro.and Mol. Biol.Rev. 70, 1032–1060.

Gaspar, I., Wippich, F., and Ephrussi, A. (2017). Enzymatic production of single-molecule FISH and RNA capture probes RNA 23,1582–1591.

George, C.X., Ramaswami, G., Li, J.B., and Samuel, C.E. (2016). Editing of Cellular Self-RNAs by Adenosine Deaminase ADAR1 Suppresses Innate Immune Stress Responses. J. Biol. Chem. 291, 6158–6168.

Gilbertson, S., Federspiel, J.D., Hartenian, E., Cristea, I.M., Glaunsinger, B. (2018). Changes in mRNA abundance drive shuttling of RNA binding proteins, linking cytoplasmic RNA degradation to transcription. Elife 7, e37663.

Glaunsinger, B., and Ganem, D. (2004). Lytic KSHV infection inhibits host gene expression by accelerating global mRNA turnover. Mol. Cell. 13, 713–23.

Gray, N.K., Hrabálková, L., Scanlon, J.P., and Smith, R.W. (2015). Poly(A)-binding proteins and mRNA localization: who rules the roost? Biochem. Soc. Trans. 43,1277–84.

Grivennikov, S.I., Greten, F.R., and Karin, M. (2010). Immunity, Inflammation, and Cancer. Cell, 140, 883–899.

Han, J. Q., Townsend, H. L., Jha, B. K., Paranjape, J. M., Silverman, R. H., and Barton, D. J. (2007). A phylogenetically conserved RNA structure in the poliovirus open reading frame inhibits the antiviral endoribonuclease RNase L. J. Virol. 81, 5561–72.

Iordanov, M. S., Paranjape, J. M., Zhou, A., Wong, J., Williams, B. R. G., Meurs, E. F., Silverman R.H., and Magun, B. E. (2000). Activation of p38 Mitogen-Activated Protein Kinase and c-Jun NH_2_-Terminal Kinase by Double-Stranded RNA and Encephalomyocarditis Virus: Involvement of RNase L, Protein Kinase R, and Alternative Pathways. Mol. Cell Biol. 20, 617–627.

Ivashki, L. B., and Donlin, L. T. (2014). Regulation of type I interferon responses. Nat. Rev. Immunol. 14, 36–49.

Jain, S., Wheeler, J.R., Walters, R.W., Agrawal, A., Barsic, A., and Parker, R. (2016). ATPase-Modulated Stress Granules Contain a Diverse Proteome and Substructure. Cell 164, 487–498.

Jensen, S., and Thomsen, A.R. (2012). Sensing of RNA Viruses: a Review of Innate Immune Receptors Involved in Recognizing RNA Virus Invasion. J. Virol. 86, 2900–2910.

Kang, J.S., Hwang, Y.S., Kim, L.K., Lee, S., Lee, W.B., Kim-Ha, J., and Kim, Y.J. (2018). OASL1 Traps Viral RNAs in Stress Granules to Promote Antiviral Responses. Mol. Cells, 41, 214–223.

Kim, D., Pertea, G., Trapnell, C., Pimentel, H., Kelley, R., and Salzberg, S.L. (2013). TopHat2: accurate alignment of transcriptomes in the presence of insertions, deletions and gene fusions. Genome Biol. 14,R36.

Kim, T. W., Hong, S., Lin, Y., Murat, E., Joo, H., Kim, T., Pascual, V., … Liu, Y. J. (2016). Transcriptional Repression of IFN Regulatory Factor 7 by MYC Is Critical for Type I IFN Production in Human Plasmacytoid Dendritic Cells. J. of Immunol. 197, 3348–3359.

Kinsella, R. J., Kähäri, A., Haider, S., Zamora, J., Proctor, G., Spudich, G., Almeida-King, J., Staines, D., Derwent, P., Kerhornou, A., et al. (2011). Ensembl BioMarts: a hub for data retrieval across taxonomic space. J. Biological Databases and Curation, bar030.

Khabar, K.S., Siddiqui, Y.M., al-Zoghaibi, F., al-Haj, L., Dhalla, M., Zhou, A., Dong, B., Whitmore, M., Paranjape, J., Al-Ahdal, M.N. et al. (2003). RNase L mediates transient control of the interferon response through modulation of the double-stranded RNA-dependent protein kinase PKR.J Biol Chem. 278, 20124–32.

Khong, A., Matheny, T., Jain, S., Mitchell, S.F., Wheeler, J.R., and Parker, R. (2017). The stress granule transcriptome reveals principles of mRNA accumulation in stress granules. Mol.Cell 68, 808–820.

Kojima, E., Takeuchi, A., Haneda, M., Yagi, A., Hasegawa, T., Yamaki, K., Takeda, K., Akira, S., Shimokata, K., and Isobe, K. (2003). The function of GADD34 is a recovery from a shutoff of protein synthesis induced by ER stress: elucidation by GADD34-deficient mice. FASEB J. 17, 1573–5.

Krug, L., Chatterjee, N., Borges-Monroy, R., Hearn, S., Liao, W.-W., Morrill, K., Prazak, L., Rozhkov, N., Theodorou, D., Hammell, M., et al. (2017). Retrotransposon activation contributes to neurodegeneration in a Drosophila TDP-43 model of ALS. PLoS Genet. 13, e1006635.

Kumar, G. R., and Glaunsinger, B. A. (2010). Nuclear Import of Cytoplasmic Poly(A) Binding Protein Restricts Gene Expression via Hyperadenylation and Nuclear Retention of mRNA. Mol. Cell Biol. 30, 4996–5008.

Langmead, B., and Salzberg, S. L. (2012). Fast gapped-read alignment with Bowtie 2. Nature methods 9, 357–9.

Li, Y., Banerjee, S., Goldstein, S.A., Dong, B., Gaughan, C., Rath, S., Donovan, J., Korennykh, A., Silverman, R.H., et al. (2017). Ribonuclease L mediates the cell-lethal phenotype of doublestranded RNA editing enzyme ADAR1 deficiency in a human cell line. eLife. 6, e25687.

Liddicoat, B.J., Piskol, R., Chalk, A.M., Ramaswami, G., Higuchi, M., Hartner, J.C., Li, J.B., Seeburg, P.H., and Walkley, C.R. (2015). RNA editing by ADAR1 prevents MDA5 sensing of endogenous dsRNA as nonself. Science 349, 1115–1120.

Lloyd, R.E. (2013). Regulation of stress granules and P-bodies during RNA virus infection. Wiley interdisciplinary reviews. RNA 4, 317–31.

Markmiller, S., Soltanieh, S., Server, K.L., Mak, R., Jin, W., Fang, M.Y., Luo, E.C., Krach, F., Yang, D., Sen, A., et al. (2018). Context-Dependent and Disease-Specific Diversity in Protein Interactions within Stress Granules. Cell 172, 590–604.

Mi, H., Huang, X., Muruganujan, A., Tang, H., Mills, C., Kang, D., and Thomas, P. D. (2016). PANTHER version 11: expanded annotation data from Gene Ontology and Reactome pathways, and data analysis tool enhancements. Nucleic Acids Res. 45, D183–D189.

Montero, H., Rojas, M., Arias, C. F., and López, S. (2008). Rotavirus Infection Induces the Phosphorylation of eIF2α but Prevents the Formation of Stress Granules. J. Virol. 82, 1496–1504.

Moon, S.L., and Parker, R. (2018). EIF2B2 mutations in vanishing white matter disease hypersuppress translation and delay recovery during the integrated stress response. RNA 24, 841–852.

Onomoto, K., Jogi, M., Yoo, J.-S., Narita, R., Morimoto, S., Takemura, A., Sambhara, S., Kawaguchi, A., Osari, S.,Nagata, K., et al. (2012). Critical Role of an Antiviral Stress Granule Containing RIG-I and PKR in Viral Detection and Innate Immunity. PLoS ONE, 7, e43031.

Pestal, K., Funk, C.C., Snyder, J.M., Price, N.D., Treuting, P.M., and Stetson, D.B. (2015). Isoforms of the RNA editing enzyme ADAR1 independently control nucleic acid sensor MDA5-driven autoimmunity and multi-organ development. Immunity 43, 933–944.

Protter, D. S. W., and Parker, R. (2016). Principles and Properties of Stress granules. Trends Cell Biol, 26, 668–679.

Rath, S., Prangley, E., Donovan, J., Demarest, K., Wingreen, N., Meir, Y., and Korennykh, A. (2018). 2-5A-Mediated mRNA Decay and Transcription Act in Concert to Reprogram Protein Synthesis during dsRNA Response. Submitted.

Reineke, L. C., Dougherty, J. D., Pierre, P., and Lloyd, R. E. (2012). Large G3BP-induced granules trigger eIF2α phosphorylation. Mol. Biol. Cell 23, 3499–3510.

Rusinova, I., Forster, S., Yu, S., Kannan, A., Masse, M., Cumming, H., Chapman, R., and Hertzog, P. J. (2012). Interferome v2.0: an updated database of annotated interferon-regulated genes. Nucleic Acids Res. 41, D1040–6.

Saldi, T.K., Ash, P.E., Wilson, G., Gonzales, P., Garrido-Lecca, A., Roberts, C.M., Dostal, V., Gendron, T.F., Stein, L.D., Blumenthal, T., et al. (2014). TDP-1, the Caenorhabditis elegans ortholog of TDP-43, limits the accumulation of double-stranded RNA. EMBO J. 33, 2947–2966.

Savan, R. (2014). Post-transcriptional regulation of interferons and their signaling pathways. J. Interferon Cytokine Res. 34, 318–29.

Schindelin, J., Arganda-Carreras, I., Frise, E., Kaynig, V., Longair, M., Pietzsch, T., Preibisch, S., Rueden, C., Saalfeld, S., et al. (2012). Fiji: an open-source platform for biological-image analysis. Nat. Methods 9, 676–82.

Schmidt, E.K., Clavarino, G., Ceppi, M., and Pierre, P. (2009). SUnSET, a nonradioactive method to monitor protein synthesis. Nat. Methods 6, 275–7.

Sidrauski, C., McGeachy, A. M., Ingolia, N. T., and Walter, P. (2015). The small molecule ISRIB reverses the effects of eIF2α phosphorylation on translation and stress granule assembly. eLife 4, e05033.

Tani, H., Mizutani, R., Salam, K. A., Tano, K., Ijiri, K., Wakamatsu, A., Isogai, T., Suzuki, Y., and Akimitsu, N. (2012). Genome-wide determination of RNA stability reveals hundreds of short-lived noncoding transcripts in mammals. Genome res., 22, 947–56.

Townsend, H. L., Jha, B. K., Silverman, R. H., and Barton, D. J. (2008). A putative loop E motif and an H-H kissing loop interaction are conserved and functional features in a group C enterovirus RNA that inhibits ribonuclease L. RNA Biol. 5, 263–72.

Trapnell, C., Hendrickson, D.G., Sauvageau, M., Goff, L., Rinn, J.L. and Pachter, L. (2013). Differential analysis of gene regulation at transcript resolution with RNA-seq. Nature Biotech. 31, 46–53.

Van Treek, B., Protter, D.S.W., Matheny, T., Khong, A., Link, C.D., and Parker, R. (2018). RNA selfassembly contributes to stress granule formation and defining the stress granule transcriptome.Proc. Natl. Acad. Sci. U.S.A. 115, 2734–2739.

Waldner, H. (2009). The role of innate immune responses in autoimmune disease development. Autoimmun, Rev. 8, 400–4.

Wreschner, D.H., James, T.C., Silverman, R.H., Kerr, I.M. (1981). Ribosomal RNA cleavage, nuclease activation and 2-5A(ppp(A2’p)nA) in interferon-treated cells.Nucleic Acids Res. 9, 1571–81.

Yang, Y. C., Di, C., Hu, B., Zhou, M., Liu, Y., Song, N., Li, Y., Umetsu, J., and Lu, Z. J. (2015). CLIPdb: a CLIP-seq database for protein-RNA interactions. BMC genomics 16, 51.

Yoo, J.-S., Takahasi, K., Ng, C. S., Ouda, R., Onomoto, K., Yoneyama, M., Lai, J.C., Lattmann, S., Nagamine, Y., Matsui, T. (2014). DHX36 Enhances RIG-I Signaling by Facilitating PKR-Mediated Antiviral Stress Granule Formation. PLoS Path.10, e1004012.

Youn, J.Y., Dunham, W.H., Hong, S.J., Knight, J.D.R., Bashkurov, M., Chen, G.I., Bagci, H., Rathod, B., MacLeod, G., Eng, S.W.M., et al. (2018). High-Density Proximity Mapping Reveals the Subcellular Organization of mRNA-Associated Granules and Bodies. Mol. Cell. 69, 517–532.

Zhang, R., Mehla, R., and Chauhan, A. (2010). Perturbation of Host Nuclear Membrane Component RanBP2 Impairs the Nuclear Import of Human Immunodeficiency Virus-1 Preintegration Complex (DNA). PLoS ONE 5, e15620.

Zhou, A., Paranjape, J., Brown, T.L., Nie, H., Naik, S., Dong, B., Chang, A., Trapp, B., Fairchild, R., Colmenares, C., and Silverman, R.H. Interferon action and apoptosis are defective in mice devoid of 2’,5’-oligoadenylate-dependent RNase L. Embo J. 1997;16:6355–63.

