## Supplementary material for "RNase L reprograms translation by widespread mRNA turnover escaped by antiviral mRNAs"

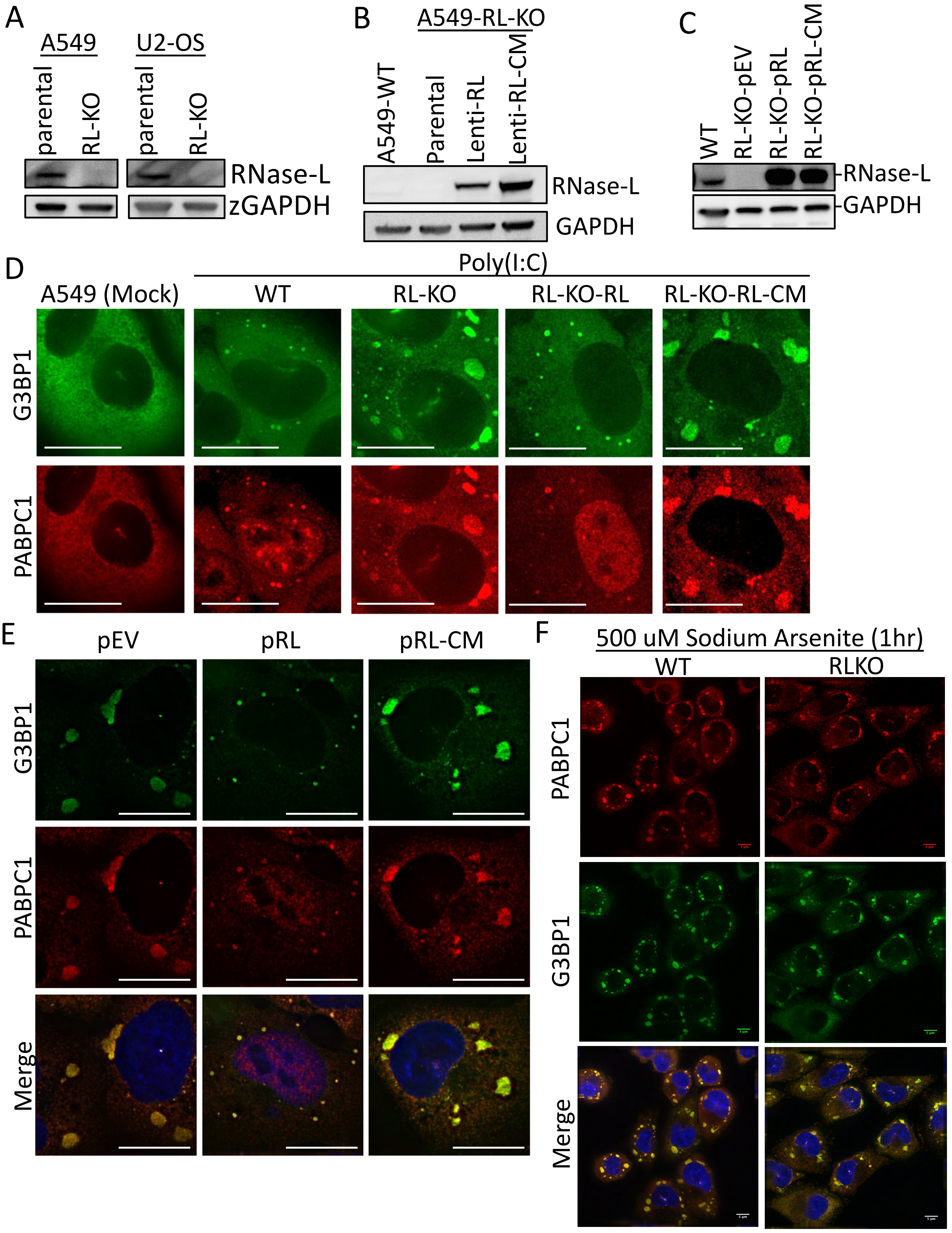


Supplemental Figure S1

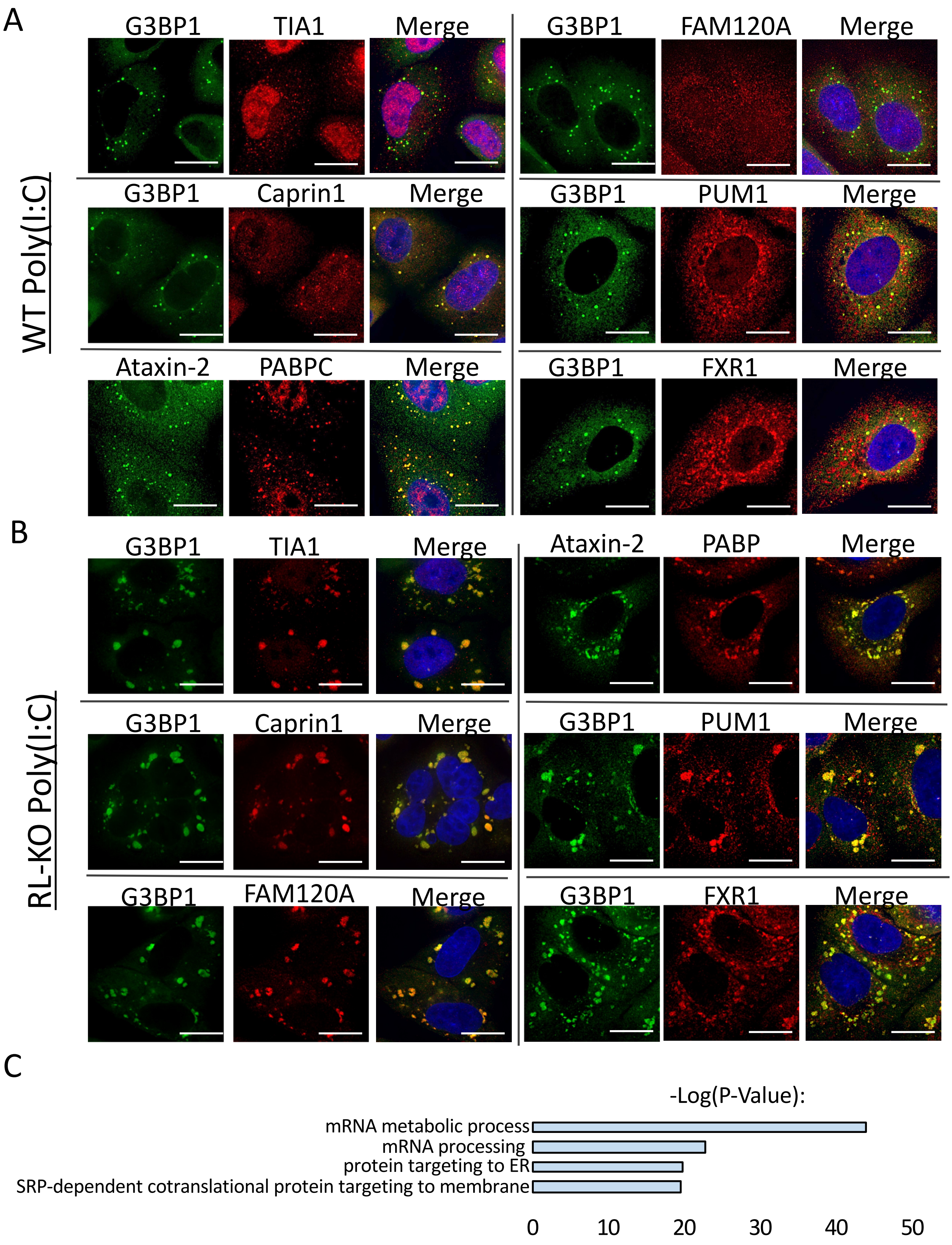


Supplemental Figure S2

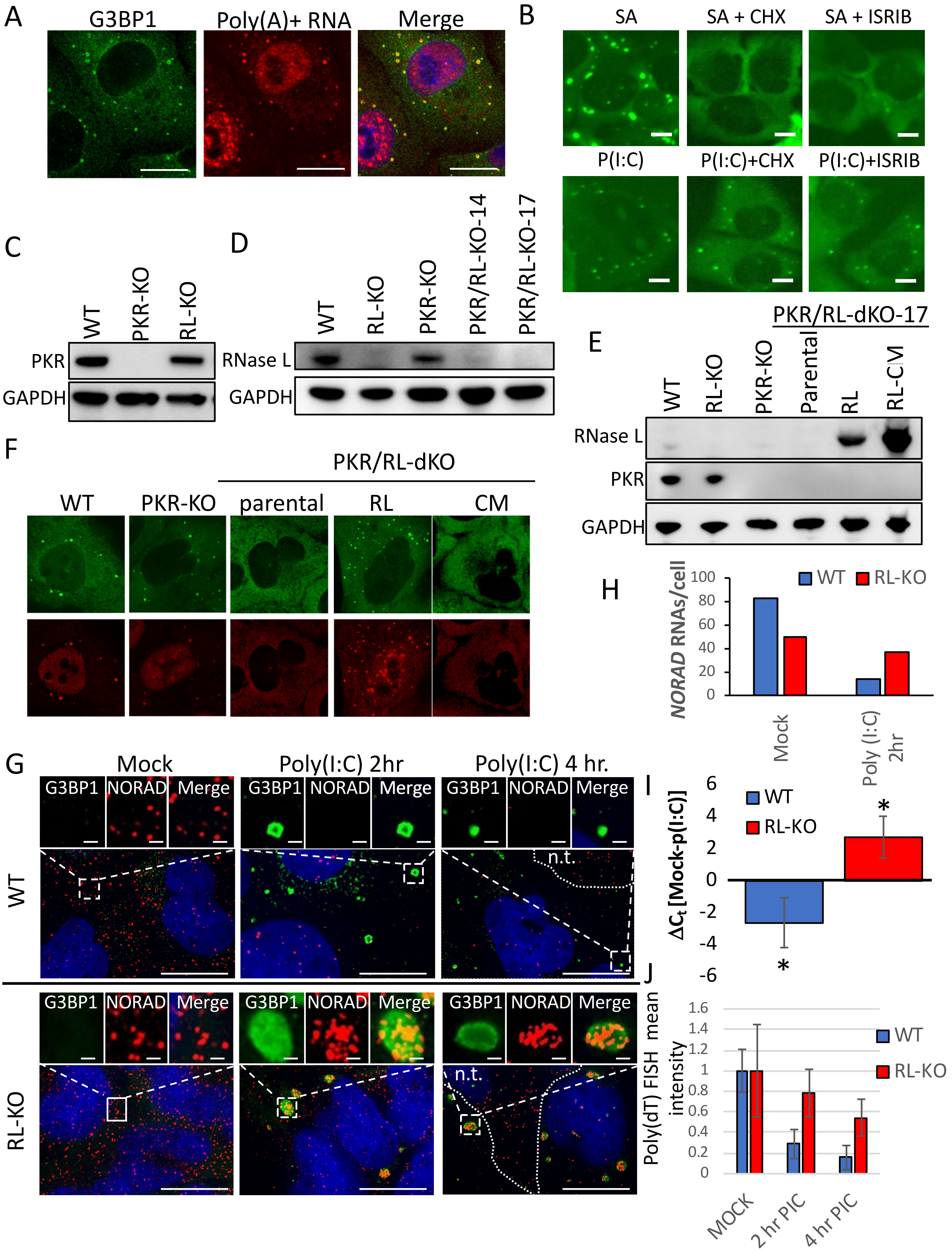


Supplemental Figure S3

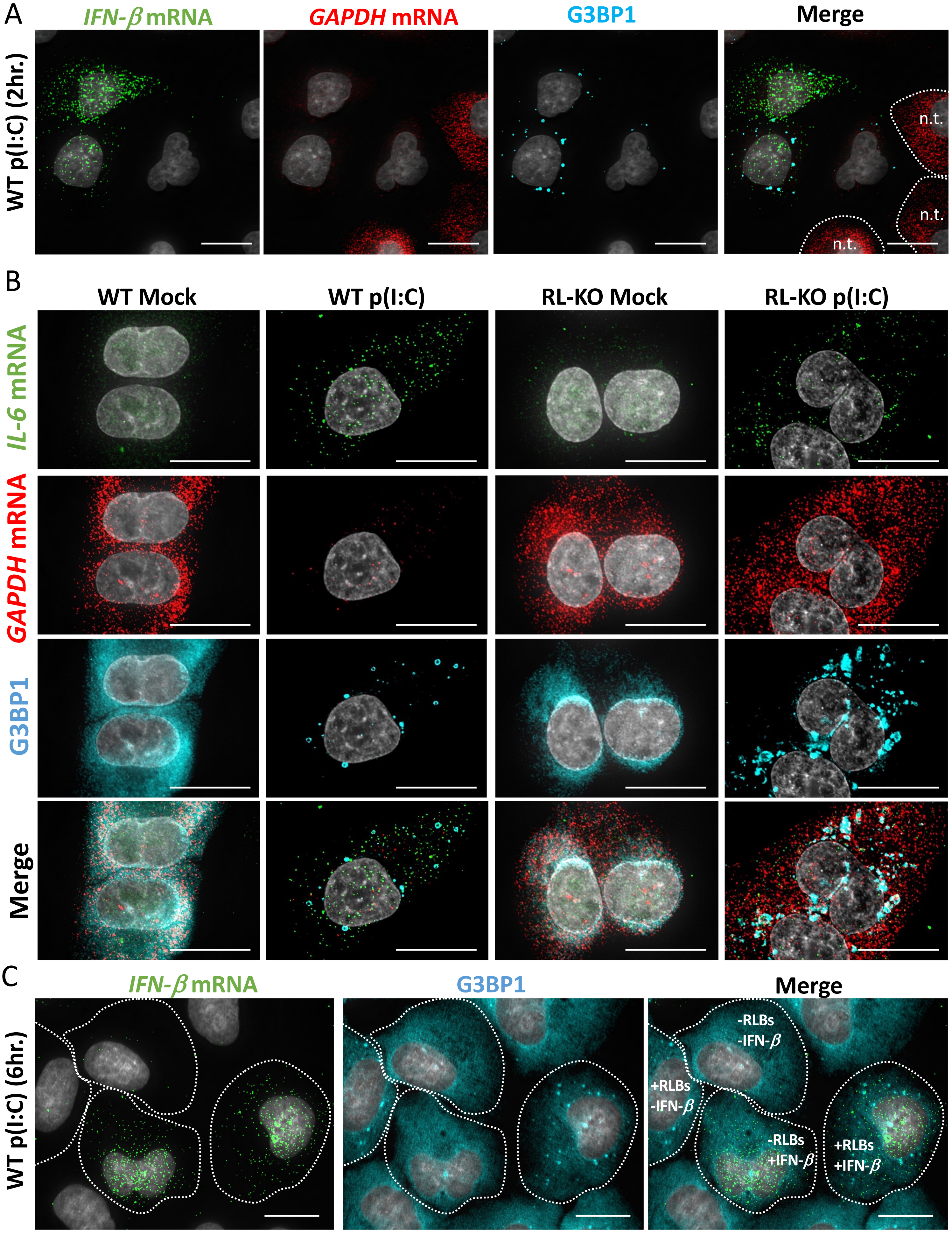


Supplemental Figure S4

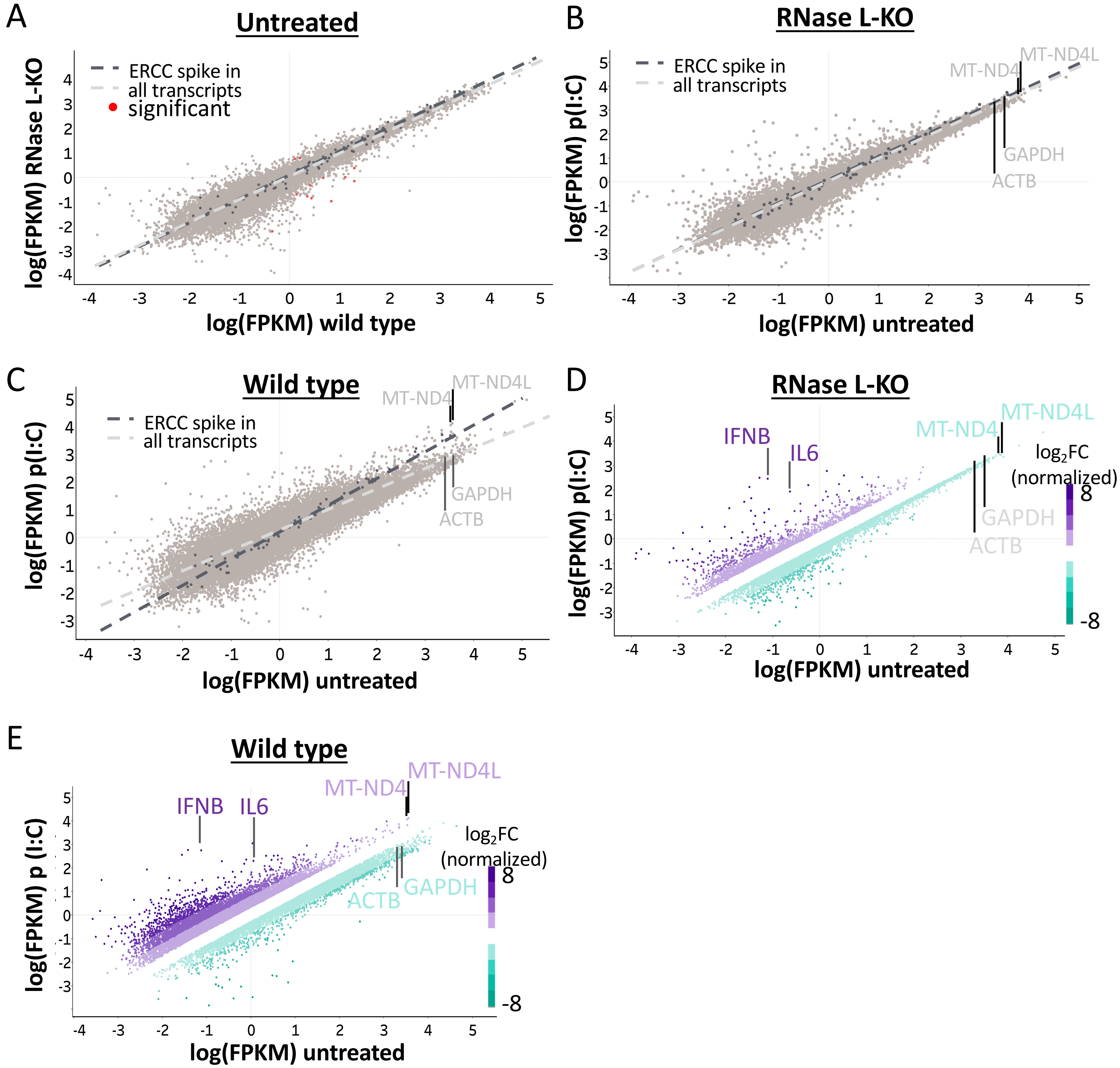


Supplemental Figure S5

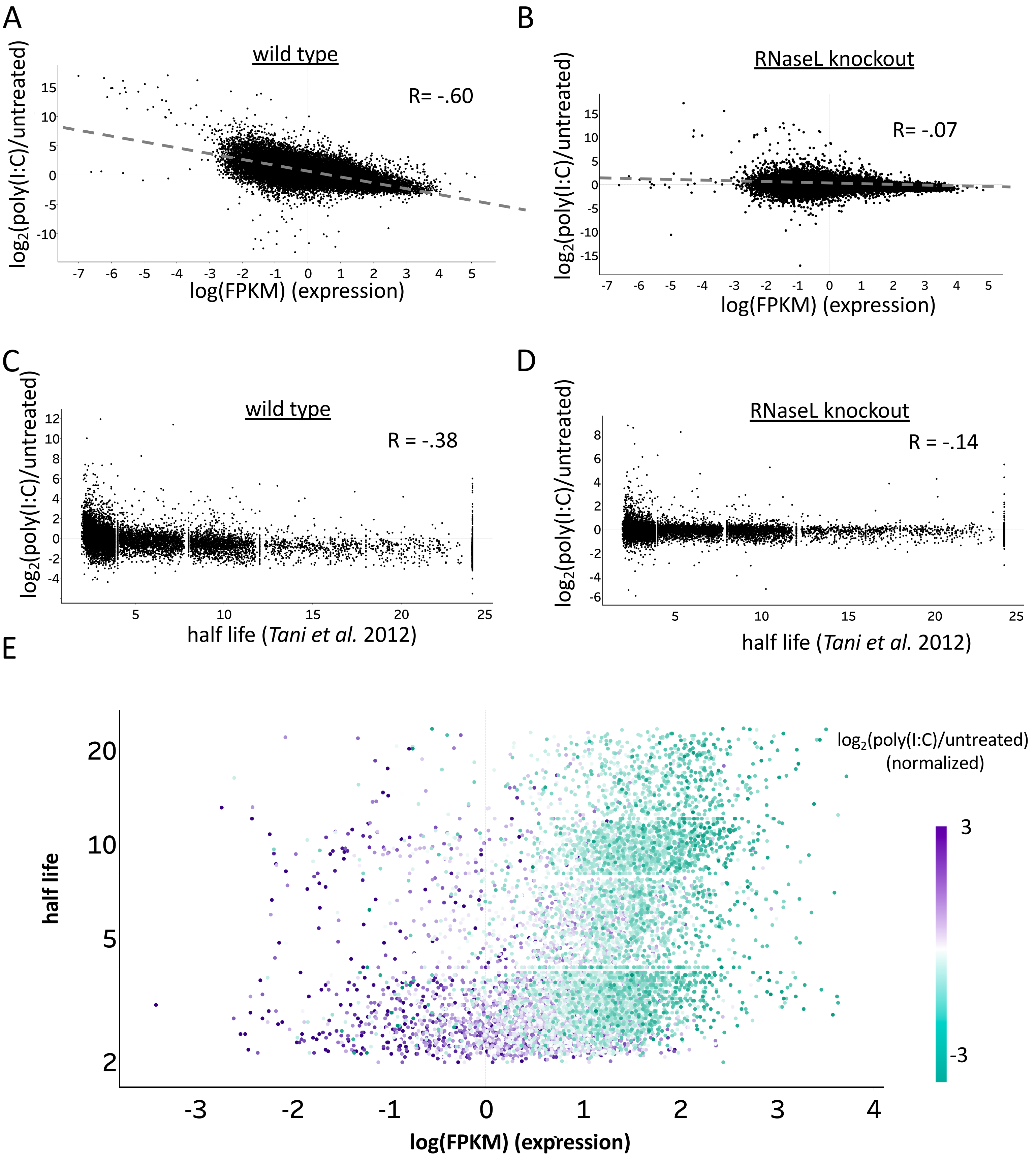


Supplemental Figure S6

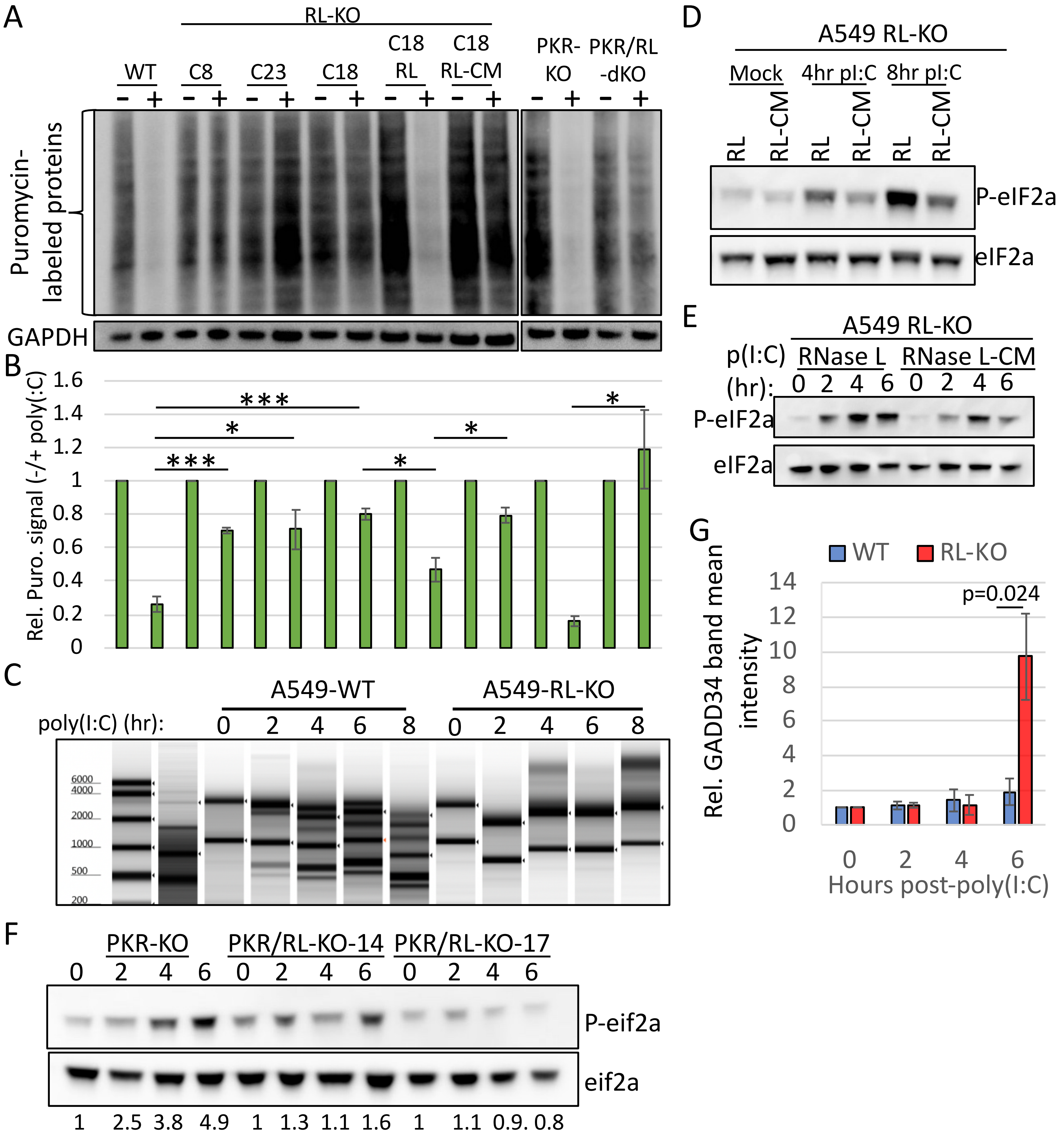


Supplemental Figure S7
